# A Non-conventional Archaeal Fluorinase Identified by *In Silico* Mining Catalyzes the Fastest Natural Nucleophilic Fluorination

**DOI:** 10.1101/2022.03.08.483453

**Authors:** Isabel Pardo, David Bednar, Patricia Calero, Daniel C. Volke, Jiří Damborský, Pablo I. Nikel

**Author notes:** Department of Microbial and Plant Biotechnology, Centro de Investigaciones Biológicas Margarita Salas, CSIC, Madrid, Spain.

## Abstract

Fluorinases, the only enzymes known to catalyze the transfer of fluorine to an organic molecule, are essential biocatalysts for the sustainable synthesis of valuable organofluorines. However, the few fluorinases identified so far have low turnover rates that hamper biotechnological applications. Here, we isolated and characterized putative fluorinases retrieved from systematic *in silico* mining and identified a non-conventional archaeal enzyme from *Methanosaeta* sp. that mediates the highest fluorination rate reported to date. Furthermore, we demonstrate enhanced production of fluoronucleotides *in vivo* in a bacterial host engineered with this archaeal fluorinase, paving the way towards synthetic metabolism for efficient biohalogenations.

Fastest biological S_N_2 fluorination

## INTRODUCTION

Fluorinated organic compounds (organofluorines), containing at least one fluorine (F) atom, are chemicals of enormous industrial interest—as evidenced by their increasing prevalence in newly developed pharmaceuticals and agrochemicals (Harsanyi and Sandford, 2015; Inoue *et al*., 2020; Ogawa *et al*., 2020). The unique physicochemical properties of F endow organofluorines with advantageous properties with respect to their non-fluorinated counterparts, e.g. increased chemical stability or improved bioavailability (O’Hagan, 2008). However, the abundance of human-made organofluorines contrasts with their relative scarcity in Nature (Carvalho and Oliveira, 2017). 5’-Fluoro-5’-deoxyadenosine (5’-FDA) synthase, or fluorinase (FlA), is the only one enzyme known to naturally catalyze the formation of the high-energy C–F bond. This enzyme, originally identified in *Streptomyces cattleya* (O’Hagan *et al*., 2002; Schaffrath *et al*., 2003; Dong *et al*., 2004), catalyzes the S_N_2 transfer of a fluoride ion (F–) to the C5’ of the essential methyl donor *S*-adenosyl-L-methionine (SAM). As indicated in step I of **Fig. 1**, this reaction generates 5’-FDA and L-methionine (L-Met) as products (Zhu *et al*., 2007). Since the discovery of FlA in 2003, only six other fluorinases have been reported in the literature, all of them sourced from Actinomycetes (Deng *et al*., 2014; Ma *et al*., 2016; Sooklal *et al*., 2020). Additionally, a chlorinase closely related to FlAs, catalyzing 5’-chloro-5’-deoxyadenosine (5’-ClDA) synthesis (step II in **Fig. 1**), has also been identified in the marine Actinomycete *Salinispora tropica* (Eustáquio et al., 2008). FlA from *S. cattleya* is capable of catalyzing the chlorination reaction as well, albeit much less efficiently than fluorination (Deng *et al*., 2006). Conversely, the SalL chlorinase cannot catalyze the formation of C–F bonds. This activity difference has been attributed to a 23-residue loop, present in all known FlAs but absent in SalL. It was hypothesized that this loop, located near the catalytic site, could influence halide specificity by modifying the architecture of the binding pocket.

**Fig. 1.**
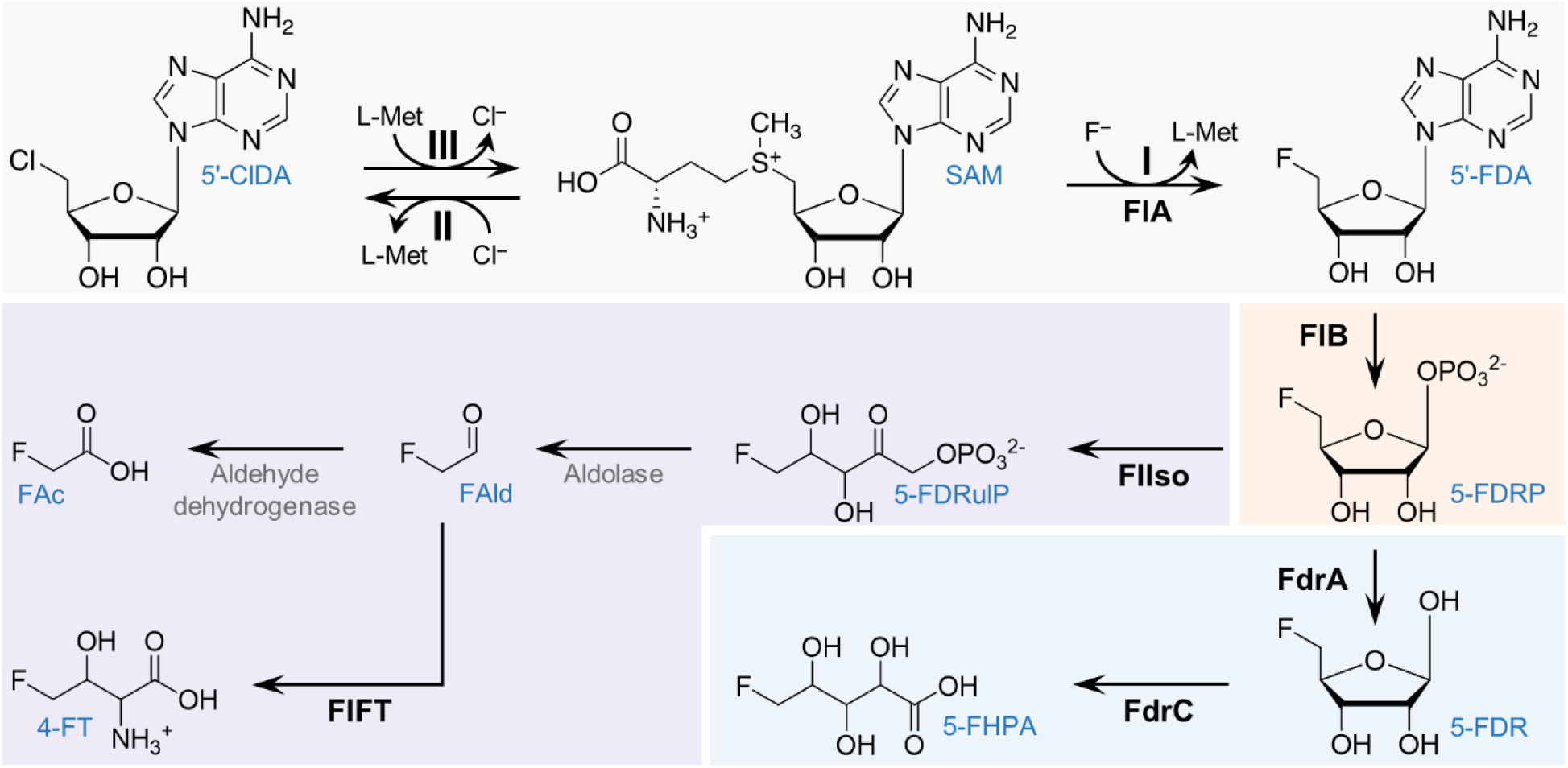
In gray, reactions catalyzed by the fluorinase/chlorinase enzyme: (I) forward fluorination reaction; (II) forward chlorination reaction; and (III) reverse chlorination reaction. In orange, common step in fluorometabolite biosynthetic pathways. In purple, canonical fluoroacetate and 4-fluoro-L-threonine biosynthetic pathway. In blue, 5’-fluoro-5’-deoxy-D-ribose biosynthetic pathway. Compound abbreviations (blue font): 5’-ClDA, 5’-chloro-5’-deoxyadenosine; SAM, *S*-adenosyl-L-methionine; 5’-FDA, 5’-fluoro-5’-deoxyadenosine; 5-FDRP, 5-fluoro-5-deoxy-D-ribose-1-phosphate; 5-FDRulP, 5-fluoro-5-deoxy-D-ribulose-1-phosphate; FAld, fluoroacetaldehyde; FAc, fluoroacetate; 4-FT, 4-fluoro-L-threonine; 5-FDR, 5-fluoro-5-deoxy-D-ribose; 5-FHPA, 5-fluoro-2,3,4-trihidroxypentanoic acid. Enzyme abbreviations (in bold): FlA, fluorinase; FlB, 5’-fluoro-5’-deoxyadenosine phosphorylase; FlIso, 5-fluoro-5-deoxy-D-ribose-1-phosphate isomerase; FlFT, 4-fluoro-L-threonine transaldolase; FdrA, 5-fluoro-5-deoxy-D-ribose-1-phosphate phosphoesterase; FdrC, 5-fluoro-5-deoxy-D-ribose dehydrogenase.

Considering the environmentally-harsh conditions currently required for the chemical synthesis of organofluorines (Cros *et al*., 2022), FlAs are promising biocatalysts for “green” production of new-to-Nature, bio-derived organofluorines and the implementation of synthetic metabolism with fluorinated intermediates in living cells (Martinelli and Nikel, 2019; Nieto-Domínguez and Nikel, 2020). However, all known FlAs are poor biocatalysts, with turnover rates < 1 min^−1^. So far, the handful of protein engineering efforts aimed at the improvement of FlA activity have had limited success (Thomsen *et al*., 2013; Sun *et al*., 2016). Furthermore, these studies mostly relied on employing surrogate substrates, e.g. 5’-ClDA, to select for enzyme variants with improved transhalogenation activity (steps III and I in **Fig. 1**). This strategy hampers the wide applicability of FlAs in a consolidated, whole-cell bioprocess where only F^−^ and an appropriate carbon substrate would be supplied as feed-stock (Calero *et al*., 2020; Markakis *et al*., 2020).

Genome-wide databases are a rich source of potentially valuable enzymes, yet their continuous, exponential expansion makes the selection of catalytically-attractive candidates challenging. The EnzymeMiner platform has been recently developed to address this issue (Hon *et al*., 2020), as an inter-active and user-friendly website (https://loschmidt.chemi.muni.cz/enzymeminer). This bioinformatic tool searches through databases upon submitting a sequence of at least one representative member of the target enzyme family, together with the identification of essential (i.e. catalytic) residues. EnzymeMiner conducts multiple database searches and accompanying calculations, which provide a set of hits and their systematic annotation based on protein solubility, possible extremophilicity, domain structures and other structural information. These collected and calculated annotations provide users with key information needed for the selection of the most promising sequences for gene synthesis, small-scale protein expression, purification, and functional characterization (Vanacek *et al*., 2018).

With the goal of expanding the FlA toolset for the biological production of organofluorines, here we describe the systematic screening and *in vitro* characterization of hitherto unknown FlAs retrieved from genome databases using EnzymeMiner. This approach led to the isolation of a non-conventional fluorinase enzyme (identified as a chlorinase) that catalyzes the fastest S_N_2 fluorination reaction reported thus far. Likewise, the implementation of this enzyme in engineered bacterial cell factories gave rise to the highest fluorometabolite content produced *via de novo* biofluorination.

## RESULTS & DISCUSSION

In an effort to identify “Nature’s best” biocatalyst, the fluorinase from *Streptomyces* sp. MA37 (FlAMA37) was used as the query sequence (Uni-Prot W0W999), and the residues D16, Y77, S158, D210, and N215 were specified as essential based on their implication in catalysis and substrate binding (**Fig. 2a**). After curing redundant sequences, sixteen unique candidates were obtained (**Table 1** and **Fig. 2b**). Some of the retrieved amino acid sequences were found to be missing several *N*-terminal residues, which were added after manually curating the deposited genome sequences from which the fluorinase genes had been predicted (**Table S1**). Out of the sixteen sequences retrieved, five corresponded to fluorinases previously reported in the literature (thus serving as an internal quality control of the prediction routine), while nine corresponded to new putative fluorinases. Another two sequences corresponded to the chlorinase from *Salinispora tropica* CNB-440 (SalL) and a putative chlorinase from the archaea *Methanosaeta* sp. PtaU1.Bin055 (FlAPtaU1). Both of these sequences lack the 23-residue loop previously hypothesized to differentiate fluorinases from chlorinases (**Fig. 2c-e**). Notably, only four of all the retrieved sequences were not sourced from Actinobacteria. These included the putative enzymes from a *Chloroflexi* bacterium (*Chloroflexi*), *Peptococcaceae* bacterium CEB3 (Clostridia), *Thermosulforhabdus norvegica* (Deltaproteobacteria) and *Methanosaeta* sp. PtaU1.Bin055 (Methanomicrobia). Phylogenetic analysis of the 16S rRNA sequences of the fluorinase-encoding organisms gave a similar result to that obtained when using the fluorinase amino acid sequences, except that, as expected, *Salinispora tropica* groups together with the other Actinomycetes, in a clade separate from the one formed by *Streptomyces* sp. (**Table S2** and **Fig. S1**).

**Table 1.** Putative fluorinases retrieved from EnzymeMiner search using FlA^MA37^ as query.

| Name | Organism | ID (%)<sup>a</sup> |
| --- | --- | --- |
| FIA<sup>MA37</sup> | *Streptomyces* sp. MA37 | Query |
| FIA<sup>Scat</sup> | *Streptomyces cattleya* | 87.6% |
| FIA<sup>Sxin</sup> | *Streptomyces xinghaiensis* | 86.0% |
| FIA<sup>SAJ15</sup> | *Streptomyces* sp. SAJ15 | 85.0% |
| FIA<sup>N902</sup> | *Actinoplanes* sp. N902-109 | 80.7% |
| FIA<sup>Amza</sup> | *Actinopolyspora mzabensis* | 78.9% |
| FIA<sup>Abar</sup> | *Amycolatopsis bartoniae* | 79.1% |
| FIA<sup>CA12</sup> | *Amycolatopsis* sp. CA-128772 | 78.6% |
| FIA<sup>AN11</sup> | *Goodfellowiella* sp. AN110305 | 77.7% |
| FIA<sup>Nbra2</sup> | *Nocardia brasiliensis* IFM 10847 | 75.7% |
| FIA<sup>Nbra3</sup> | *Nocardia brasiliensis* NCTC 11294 | 75.3% |
| FIA<sup>Nbra1</sup> | *Nocardia brasiliensis* ATCC 700358 | 75.3% |
| FIA<sup>Cbac</sup> | *Chloroflexi* bacterium | 69.3% |
| FIA<sup>Pbac</sup> | *Peptococcaceae* bacterium CEB3 | 64.8% |
| FIA<sup>Tnor</sup> | *Thermodesulforhabdus norvegica* | 54.5% |
| FIA<sup>PtaU1</sup> | *Methanosaeta* sp. PtaU1.Bin055 | 49.5% |
| SalL<sup>Stro</sup> | *Salinispora tropica* CNB-440 | 35.6% |

<sup>a</sup> Sequence identity. References to known FIAs are indicated.

**Fig. 2.**
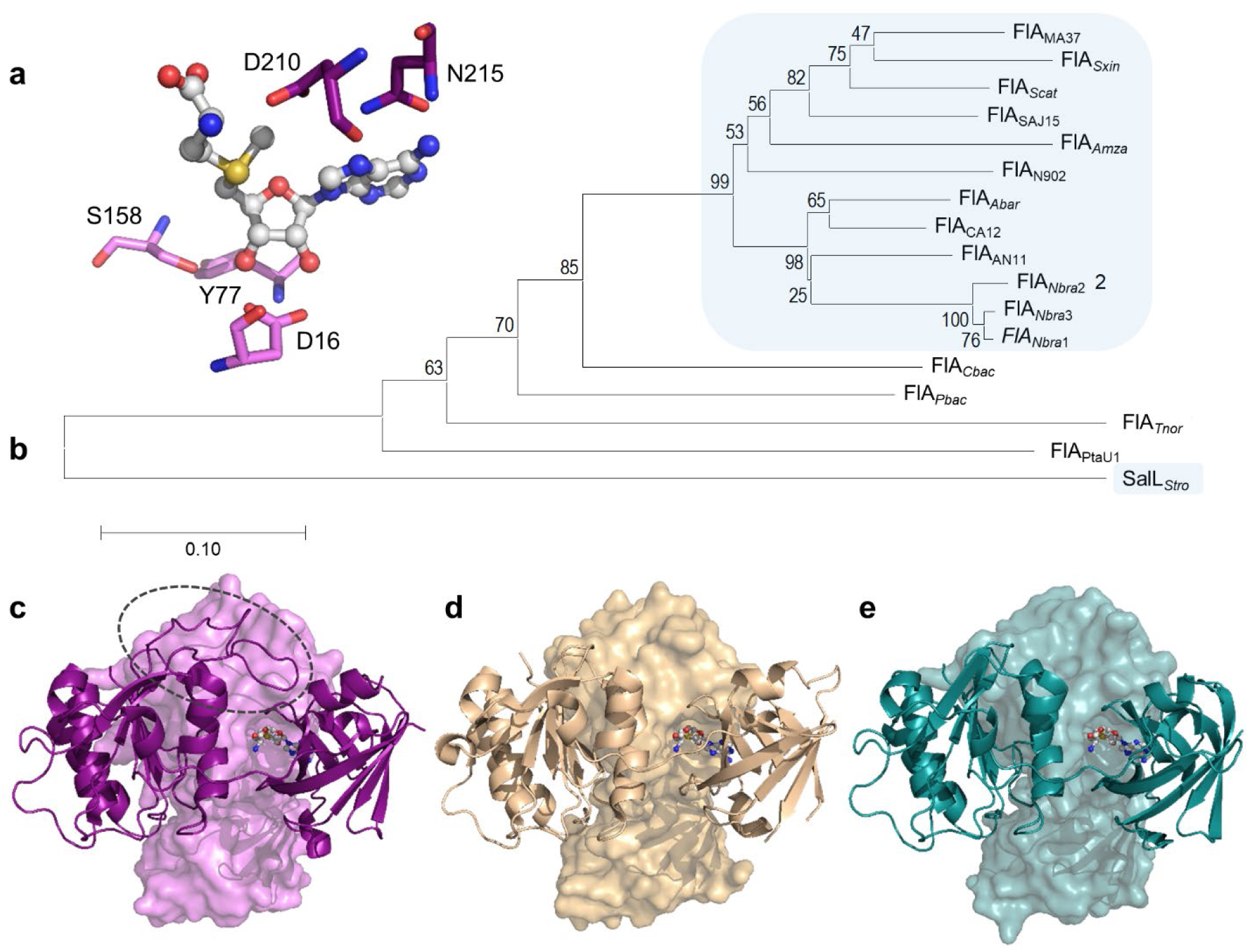
Putative fluorinases identified by genome mining. (**a**) Residues specified as essential for the EnzymeMiner search, based on the crystal structure of FlA^MA37^ (PDB ID 5B6I). The SAM substrate is shown in ball-and-stick representation. (**b**) Phylogenetic tree of retrieved fluorinase sequences obtained using MEGA-X software, inferred using the Neighbor-Joining method with a bootstrap of 10,000 iterations. The percentage of replicate trees in which the associated taxa clustered together in the bootstrap test are shown next to the branches. The tree is drawn to scale, with branch lengths in the same units as those of the evolutionary distances used to infer the phylogenetic tree. Sequences sourced from Actinomycetes are highlighted as blue squares. (**c-e**) 3D structures for FlA^MA37^ (**c**), SalL^*Stro*^ (**d**, PDB ID 6RYZ), and FlA^PtaU1^ (e, modelled from the FlA^*Scat*^ crystal structure PDB ID 2V7V). The loop hypothesized to differentiate fluorinases from chlorinases is circled in a dashed gray line. Two chains from the homo-trimer for each structure are shown as cartoon and surface representations, respectively.

The genomic context of the different *flA* genes was likewise examined (**Table S3**). As reported for the fluorination gene clusters of *Streptomyces* sp. MA37, *N. brasiliensis, Actinoplanes* sp. N902-109, and *S. xinghaiensis*, all Actinomycetes presented gene clusters resembling that of *S. cattleya* (**Fig. 3**), the most studied source of *fl* genes to date (Huang *et al*., 2006; O’Hagan and Deng, 2015). The genes *flB* (encoding a 5’-FDA phosphorylase), *flG* (encoding a response regulator), *flH* (encoding a putative cation:H^+^ antiporter) and *flI* (encoding a *S*-adenosyl-L-homocysteinase) were conserved in all Actinomycetes. Most of them also presented the genes *flIso* (5’-fluoro-5’-deoxyadenosine-1-phosphate isomerase) and *flFT* (4-fluoro-L-threonine transaldolase), involved in the synthesis of fluoroacetate and 4-fluoro-L-threonine, respectively. In these cases, genes encoding a prolyl-tRNA synthetase-associated protein and an EamA family transporter were usually found in proximity to *flFT*. In *S. cattleya*, the products of these genes (termed *fthB* and *fthC*, respectively) play a role in detoxification by deacylation of 4-fluoro-L-threoninyl·tRNA and export of 4-fluoro-L-threonine (McMurry and Chang, 2017). Interestingly, *Amycolatopsis bartoniae* and *Goodfellowiella* sp. AN110305 lacked either *flIso* and *flFT* orthologues within their *fl* clusters, presenting, instead, orthologues to the *fdr* genes of *Streptomyces* sp. MA37 that are probably involved in the biosynthesis of 5-fluoro-2,3,4-trihydroxypentanoic acid *via* the fluorosugar intermediate 5-fluoro-5-deoxy-D-ribose (Ma *et al*., 2015). Activities encoded in these clusters include phosphoesterases, short chain dehydrogenases, dihydroxyacid dehydratases and cyclases, suggesting that the main fluorinated compounds produced by these organ-isms could be different from the canonical fluoroacetate and 4-fluoro-L-threonine. Similar activities seem to be also encoded by genes in the vicinity of *flA* in *Chloroflexi* bacterium and *salL* in *S. tropica* (Eustáquio *et al*., 2009). Other genes widely distributed amongst the different actinomycotal clusters encoded activities related to SAM synthesis (i.e. SAM synthetase) and *S*-adenosyl-L-homocysteine degradation (i.e. *S*-adenosyl-L-homocysteinase), a competitive inhibitor of fluorinase activity (Schaffrath *et al*., 2003). As indicated above, the latter gene (*flI*) was present in all actinomycotal clusters. Since SAM and *S*-adenosyl-L-homocysteine are involved in essential cellular reactions, it is likely that these enzymes modulate the levels of these compounds during secondary metabolism, when organofluorines are produced. Further analysis of the genes found in these *fl* clusters will provide clues as to what activities are needed to establish robust and efficient biofluorination pathways in heterologous hosts.

**Fig. 3.**
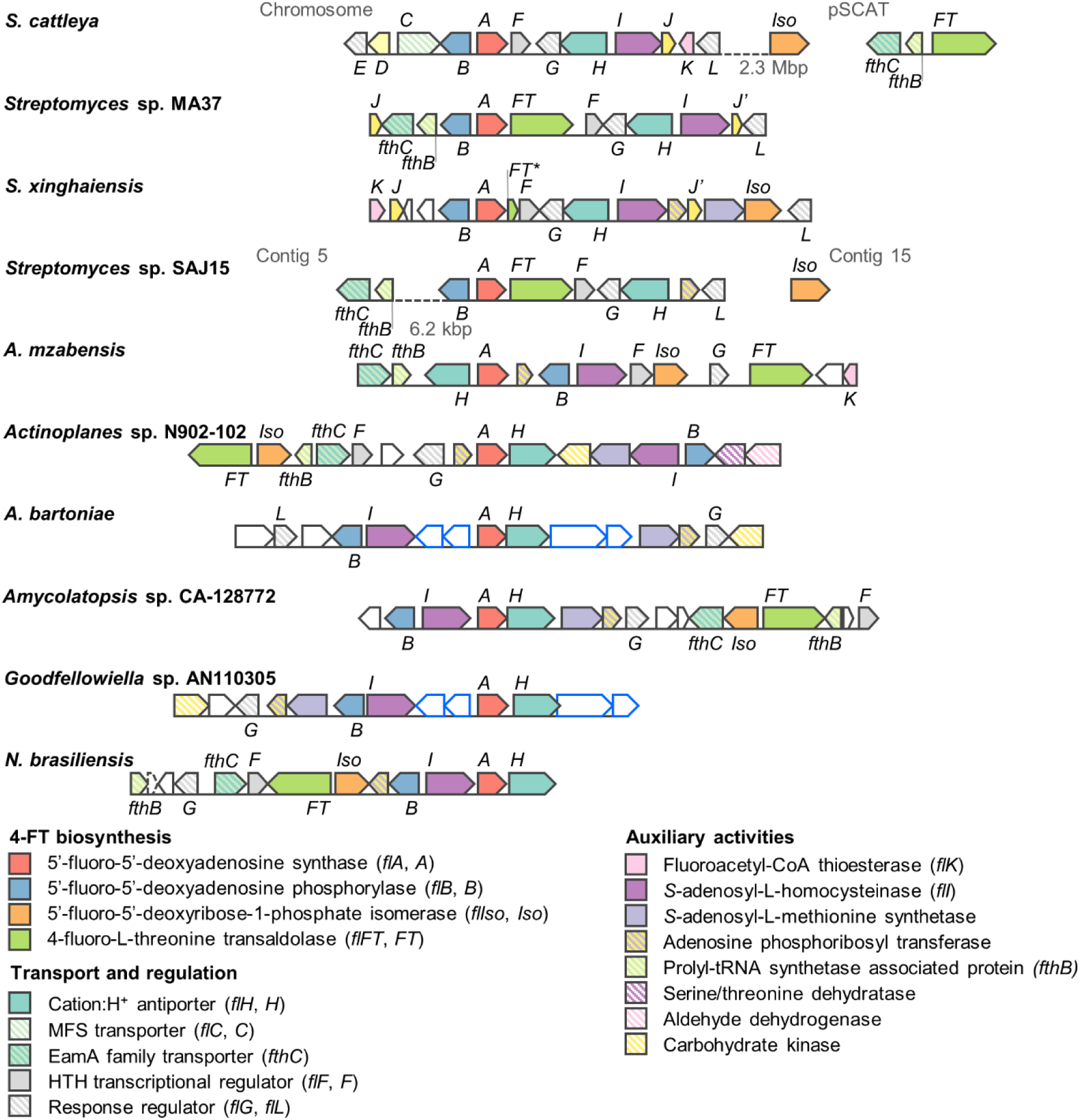
Fluorination gene clusters from Actinomycetes. For clarity, the clusters are drawn centered on *flA* (*A*) in the sense orientation. Numbers under dashed lines indicate the distance between open reading frames (ORFs) found in the same sequence entry; ORFs in separate entries are not connected by a line. Italicized letters indicate orthologues to the corresponding *fl* genes from *S. cattleya. J’* indicates duplicate *flJ* copies (encoding DUF190 domain-containing protein). *FT\** is a truncated pseudogene homologous to *flFT*. Orthologues to *fdr* genes from *Streptomyces* sp. MA37 are indicated as white blocks with blue outlines. ORFs outlined in black represent genes with other/unknown functions. MFS, major facilitator superfamily; HTH, helix-turn-helix.

Next, coding sequences of all FlA candidates were codon-optimized for production in *Escherichia coli* as *N*-terminal His-tag fusions (**Tables S4** and **S5**)—*flA*^MA37^, *flA*^*Scat*^ and *flA*^*Sxin*^ had been previously codon-optimized for expression in Gram-negative hosts (Calero *et al*., 2020). SalL^*Stro*^ was not included in this experimental set since it is reportedly inactive on F^−^ (Eustáquio *et al*., 2008). The expression of the 16 candidate genes was initially evaluated in 96-well microtiter plate cultures. FlA^*Tnor*^, FlA^*Amza*^ and FlA^*Pbac*^ could not be obtained as soluble enzymes and were not included in further analyses. Moreover, very faint bands of the expected size were observed in SDS-PAGE of *E. coli* extracts producing FlA^*Tnor*^ and FlA^*Amza*^, suggesting limited expression levels (**Fig. S2**). Therefore, we proceeded to obtain the remaining 13 candidates in medium-scale shaken-flask cultures for His-tag purification and activity assays. The purified enzymes were incubated in the presence of increasing SAM concentrations for 1 h, after which the 5’-FDA produced was measured by HPLC. 5’-FDA synthase activity could be detected for 12 out of the 13 candidates (**Fig. S3**). The protein concentration was normalized for these assays, although the enzymes were recovered with varying degrees of purity due to differences in solubility—typical of proteins isolated from high-G+C-content species when produced in a Gram-negative host. Notably, the enzyme from *Methanosaeta* sp. (FlA^PtaU1^, predicted to be a chlorinase), was one of the top performers, together with FlA^SAJ15^. These enzymes had specific activities similar to those of FlA^MA37^ and FlA^*Sxin*^, which present the highest catalytic efficiencies on SAM fluorination reported to date.

FlA^PtaU1^ and FlA^SAJ15^ were selected for large-scale shaken-flask production and a more detailed biochemical characterization. Steady-state kinetics assays with 1 μM of the purified protein, varying concentrations of SAM (1.5 to 800 μM) and 75 mM KF revealed that both of these enzymes presented higher turnover rates (k_*cat*_) than FlA^MA37^ and FlA^*Sxin*^ (**Fig. 4a** and **Table 2**). Surprisingly, K_*M*_^SAM^ values were < 10 μM, much lower than what had been previously reported in the literature for fluorinases (Schaffrath *et al*., 2003; Zhu *et al*., 2007; Sooklal *et al*., 2020). Previous studies used high enzyme concentrations (>10 μM), which impedes reaching a steady state of the reaction for substrate concentrations below 10 μM. Additionally, we have used a KF concentration that ensures F– saturation without causing any inhibitory effect (previous studies have used KF concentrations > 200 mM).

**Fig. 4.**
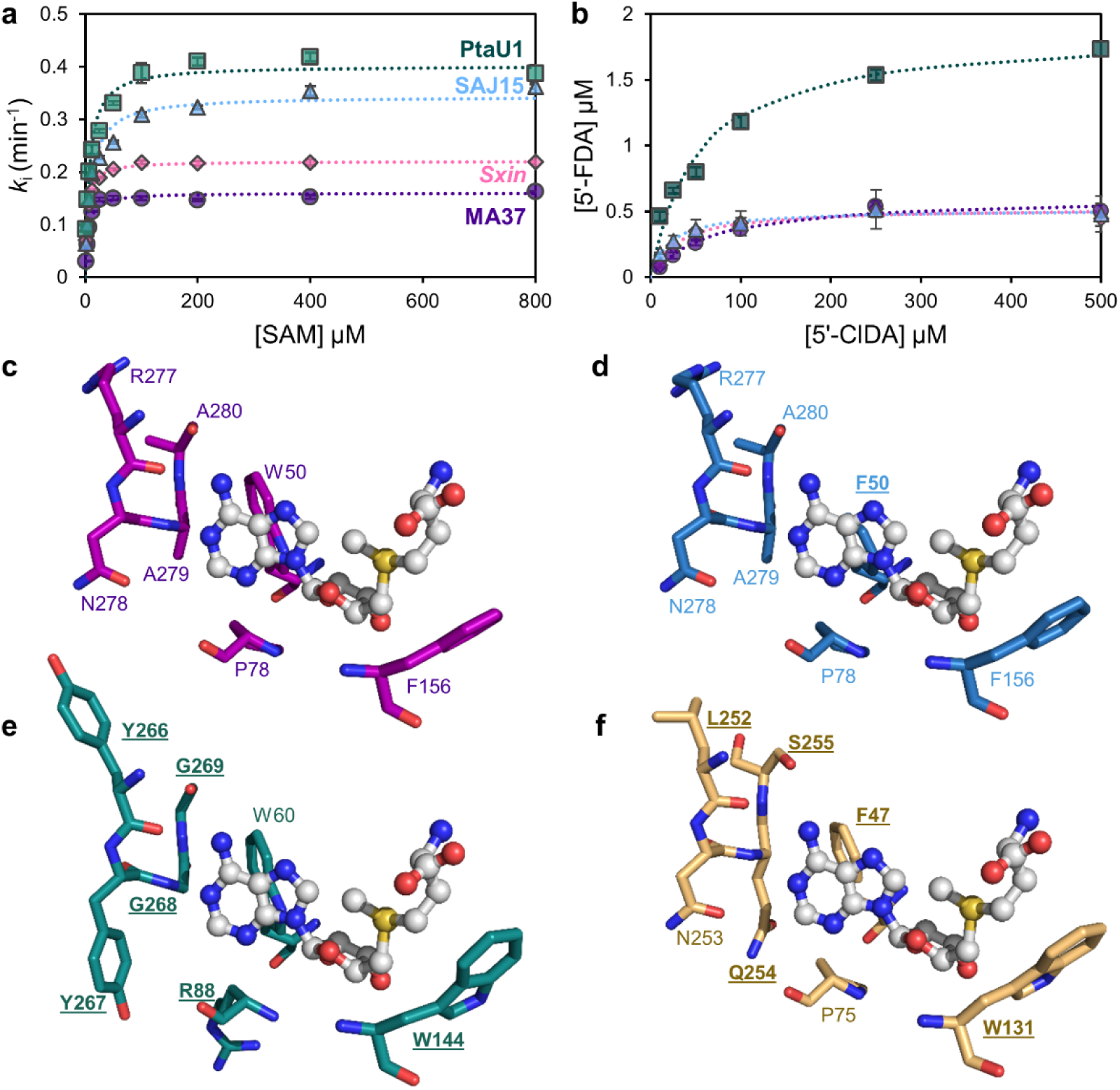
Biochemical characterization and residue conservation of selected fluorinases. (**a**) Steady-state fluorination assays using increasing SAM concentrations. Reactions were carried out at 37°C in 50 mM HEPES buffer, pH 7.8, with 75 mM KF. Dotted lines show fits to the Michaelis-Menten equation (R^2^ > 0.95 in all cases). (**b**) End-point (1 h) transhalogenation assays with increasing 5’-ClDA concentrations. Reactions were carried out at 37°C in 50 mM HEPES buffer, pH = 7.8, with 75 mM KF and 1 mM L-Met. Error bars represent standard deviations from triplicate independent assays. Symbols and color codes are kept in both panels. (**c-f**) Variable residues in the substrate binding pocket of FlA^MA37^ (**c**), FlA^SAJ15^ (**d**), FlA^PtaU1^ (**e**) and SalL^*Stro*^. Residues that differ from those of FlA^MA37^ are labelled in underlined bold font. FlA^*Sxin*^ residues are identical to those of FlA^MA37^. The SAM substrate is shown in ball-and-stick representation.

**Fig. 5.**
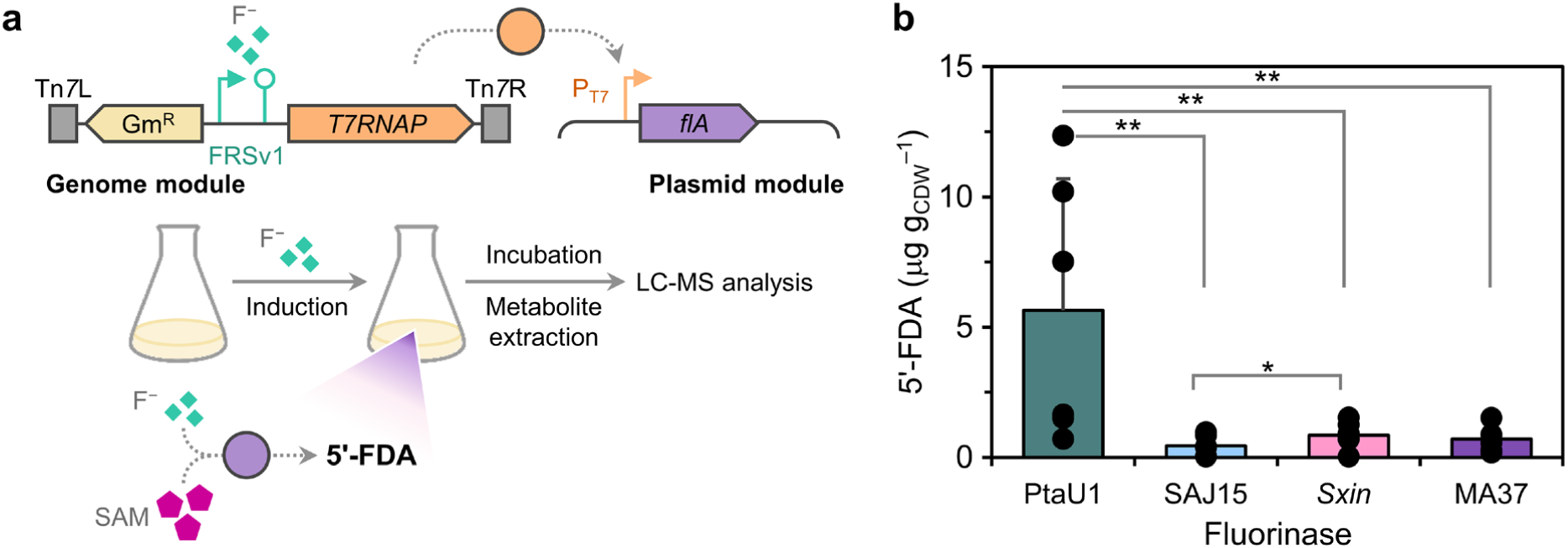
Implementing *in vivo* biofluorination in engineered *P. putida* KT2440. **(a)** Schematic representation of the fluoride-responsive genetic circuit based on the T7 phage RNA polymerase (T7RNAP) described by Calero *et al*. (2020) and workflow for the *in vivo* biofluorination assays in *P. putida*. The fluoride-responsive genetic circuit was induced by the addition of 15 mM NaF to the cultures at an OD_600_ = 0.4–0.6. After incubating the cultures at 30°C for 20 h, an aliquot was taken for metabolite extraction, biomass quantification and LC-MS analysis of fluorometabolites. Further details are provided in the Supporting Information. **(b)** Intracellular 5’-FDA content in engineered *P. putida* expressing different fluorinase genes. The fluorometabolite concentration was normalized by the cell dry weight (CDW). Black dots show individual values of independent experiments and error bars represent standard deviations. Asterisks indicate significant differences with *p*-values < 0.1 (*) or <0.05 (**) for a two-sample, one-sided Welch’s *t*-test.

**Table 2.** Michaelis-Menten kinetic constants of selected fluorinases.^*a*^

| Fluorinase | $K_M^{\text{SAM}}$<br>($\mu\text{M}$) | $k_{\text{cat}}$<br>($\text{min}^{-1}$) | $k_{\text{cat}}/K_M^{\text{SAM}}$<br>($\text{mM}^{-1} \text{min}^{-1}$) |
| --- | --- | --- | --- |
| FIA<sup>MA37</sup> | $4.42 \pm 0.58$ | $0.16 \pm 0.01$ | $36.36 \pm 4.82$ |
| FIA<sup>Sxin</sup> | $3.76 \pm 0.15$ | $0.22 \pm 0.01$ | $58.63 \pm 2.63$ |
| FIA<sup>SAJ15</sup> | $9.62 \pm 1.43$ | $0.34 \pm 0.01$ | $35.81 \pm 5.43$ |
| FIA<sup>PtaU1</sup> | $6.99 \pm 1.06$ | $0.41 \pm 0.01$ | $57.54 \pm 8.85$ |

<sup>a</sup> Assays conducted in 50 mM HEPES, pH 7.8, with 75 mM KF and varying SAM concentrations at 37°C.
Average and standard deviation are given for triplicate independent measurements.

To gain insight on the structural factors that could determine these differences in activity, we inspected the predicted crystal structures of FlA^MA37^, FlA^*Sxin*^, FlA^SAJ15^, FlA^PtaU1^ and SalL^*Stro*^. Examination of the residues potentially interacting with SAM (distances < 5 Å) revealed important variations between the substrate binding pocket of FlAPtaU1 and that of the other fluorinases (**Fig. 4c-f**). These differences were mostly located near the adenyl moiety of SAM, and involve the substitution of a conserved proline for an arginine residue and an RNAA motif for YYGG. This motif is found in the *C*-terminal domain of fluorinases, which is more variable than the *N*-terminal domain (**Fig. S4**) and is presumably also involved in hexamer formation (Kittilä *et al*., 2022). Interestingly, the variations found in FlA^PtaU1^ do not resemble those seen in the SalL^*Stro*^ chlorinase. The evaluation of the effect of these amino acid differences in fluorinase activity will be of interest for future enzyme engineering efforts.

Since FlA^PtaU1^ was predicted to be a chlorinase, we evaluated whether it was also active towards S_N_2-dependent addition of Cl– onto SAM. Unexpectedly, no 5’-ClDA accumulation could be detected in enzymatic reactions in which KF was replaced by KCl—in contrast to SalL^*Stro*^ (Eustáquio *et al*., 2008). Previous studies have shown that FlA^*Scat*^ can also catalyze the chlorination reaction (Deng *et al*., 2006). However, this feature requires the simultaneous removal of L-Met or 5’-ClDA, the reaction products, since the reverse dehalogenation reaction is favored. We could observe transhalogenation on 5’-ClDA (i.e. 5’-FDA production in the presence of L-Met and F–, steps III and I in **Fig. 1**, respectively; see **Fig. 3b**). Again, FlA^PtaU1^ catalytically outperformed all other fluorinases, with a 3-fold higher V_*max*_ value. Although we cannot rule out that FlAPtaU1 could execute *de novo* chlorination, it is clear that the 23-residue loop reportedly found in «conventional» fluorinases is not essential for the activity towards F–.

On this background, we tested the biosynthesis of fluorometabolites both *in vitro* and *in vivo* by engineering selected fluorinases in *Pseudomonas putida*. We have previously designed a fluoride-responsive genetic circuit that enabled biofluorination in this heterologous host (Calero *et al*., 2020). Here, this system was adapted to express either *flA*^PtaU1^ or *flA*^SAJ15^, codon-optimized to facilitate expression in *P. putida*. Upon induction of the system with NaF and expression of the fluorinase genes for 20 h at 30°C, production of 5’-FDA was determined by LC-MS to evaluate *de novo* fluorination activity *in vivo* (**Fig. 4**). Production of 5’-FDA could be detected for cells expressing either *flA*^PtaU1^, *flA*^SAJ15^, *flA*^MA37^ and *flA*^*Sxin*^. Notably, intracellular 5’-FDA, indicative of *in vivo* biofluorination, was 6- to 12-fold higher in cells expressing *flA*^PtaU1^ with respect to the other three fluorinase genes. On the other hand, fluorinase activity from cell-free extracts incubated for 20 h at 30°C in the presence of exogenously-added 200 μM SAM and 5 mM NaF was similar in all cases (data not shown).

## CONCLUSION

In conclusion, out of the 10 newly identified enzymes, the non-conventional FlA from the archaea *Methanosaeta* sp. PtaU1.Bin055 was found to present turnover rates far superior than those of all FlAs reported to date. Surprisingly, this enzyme lacks the loop that was hypothesized to be a differentiating feature between fluorinases and chlorinases, challenging the prevailing view that this loop is required for activity towards F–. This work highlights the importance of systematic and efficient biocatalyst selection across ever-expanding genomic databases followed by careful characterization *in vitro* and cell factory engineering *in vivo*. These results open avenues for the implementation of neo-metabolic pathways incorporating F in the host cell by synthetic biology approaches (Cros *et al*., 2022).

## ABBREVIATIONS

5’-ClDA: 5’-chloro-5’-deoxyadenosine
5’-FDA: 5’-fluoro-5’-deoxyadenosine
FlA: fluorinase
L-Met: L-methionine
SAM: *S*-adenosyl-L-methionine
SalL: chlorinase.

