## Supplemental Data 1 for "A Non-conventional Archaeal Fluorinase Identified by *In Silico* Mining Catalyzes the Fastest Natural Nucleophilic Fluorination"

*Corresponding author*

### MATERIALS AND METHODS

***In silico* screening of putative fluorinases, sequence analysis, and structure modelling.** The *EnzymeMiner* 1.0 online tool (Hon *et al.*, 2020), available online at https://loschmidt.chemi.muni.cz/enzymeminer/, was used to screen putative fluorinases, using the $FlA^{MA37}$ sequence as query (UniProt accession no. W0W999) and introducing D16, Y77, S158, D210 and N215 as the essential residues needed for catalysis. Results can be accessed through the *EnzymeMiner* webpage under Job ID 83urmz. The retrieved sequences were cured to remove duplicates and to add, when relevant, missing *N*-terminal residues. Multiple sequence alignment and phylogeny analysis were conducted with *MEGA X* software using the Neighbor-Joining method (Kumar *et al.*, 2018). Phylogenetic analysis using 16S rRNA sequences were performed essentially in the same way. Sequence analysis of the fluorination gene clusters was performed by conducting tBLASTn searches (Gertz *et al.*, 2006) against whole-genome sequences of the different organisms deposited in the National Center for Biotechnology Information (NCBI) databases, using the Fl protein sequence from *S. cattleya* (**Table S3**) and the Fdr protein sequence from *Streptomyces* sp. MA37 (GenBank accession no. LN612605.1) as query. Protein structure models were created using the SWISS-MODEL server (Waterhouse *et al.*, 2018) and visualized with PyMOL Molecular Graphics System (Schrödinger, LLC). For residue conservation visualization, the ConSurf server (Ashkenazy *et al.*, 2016) was used to estimate conservation scores and the results were visualized with the UCSF ChimeraX software (Pettersen *et al.*, 2004).

**Production of fluorinases in *Escherichia coli* and protein purification procedures.** Synthetic genes encoding the different fluorinases in this study were codon-optimized for expression in *E. coli* and synthesized as gBlocks by Integrated DNA Technologies. These fragments were then inserted into a modified pET-28a(+) vector as *N*-terminal His-tag fusions by USER cloning (**Tables S4** and **S5**). The $flA^{MA37}$, $flA^{Scat}$ and $flA^{Sxin}$ genes, previously codon-optimized for expression in Gram-negative bacteria (e.g. *P. putida*), were amplified by PCR from the corresponding plasmids and also cloned into the pET-28a(+) backbone. These modified vectors encode a TEV cleavage site between the His-tag and the protein of interest instead of the standard thrombin digestion site. Here, the native *START* codon was removed from the fluorinase genes to obtain gene fusions. Plasmid construction and maintenance was performed in *E. coli* DH5$\alpha$ $\lambda$*pir* competent cells according to standard molecular biology protocols (Green and Sambrook, 2012). Following sequence verification, the plasmids were transformed into *E. coli* BL21(DE3) for protein production. Kanamycin (Km, 50 mg/L) was added to the culture media for plasmid selection. Strain stocks were maintained at –80°C in 25% (v/v) glycerol.

Different scales were used for the production of fluorinases depending on the downstream application. In all cases, pre-cultures inoculated from glycerol stocks were incubated overnight in lysogeny broth (LB medium) + Km at 37°C with constant agitation. For 96-deep well plate production of fluorinases, overnight cultures were diluted 100-fold in 0.5 mL of 2×YT medium + Km in duplicates. Upon incubating for 2.5 h at 37°C and 250 rpm, one replicate was induced with 1 mM isopropyl $\beta$-D-1-thiogalactopyranoside (IPTG), after which the plate was incubated overnight at 16°C and 150 rpm. An aliquot was removed for SDS-PAGE analysis of total protein content; the rest of the cells were harvested by centrifugation and lysed by adding 100 $\mu$L BugBuster Master Mix (Novagen). Following ~20 min incubation at room temperature with shaking, plates were centrifuged again and the supernatants were collected for SDS-PAGE analysis of the soluble fraction.

Medium-scale shaken-flask cultivations were carried out in 250-mL baffled Erlenmeyer flasks containing 50 mL 2×YT medium + Km, and inoculated from overnight cultures to an optical density measured at 600 nm ($OD_{600}$) = 0.05. Cells were grown at 37°C and 250 rpm to $OD_{600}$ = 0.6-0.8. At this point, gene expression was induced with 1 mM IPTG, and the culture was further incubated for 24 h at 16°C and 150 rpm. Thereafter, the cells were harvested by centrifugation and cell pellets were frozen at –20°C until protein extraction. For this step, cells were resuspended in 2 mL of His A buffer (300 mM NaCl and 20 mM imidazole in 10 mM HEPES, pH = 7.5) and disrupted with glass beads in a Precellys 24 homogenizer (Bertin Instruments) with two 20 s cycles at 6,000×*g*.

In the case of large-scale shaken-flask cultivations, overnight cultures were diluted 100-fold in 50 mL 2×YT medium + Km in 250-mL baffled flasks and grown at 37°C and 250 rpm to $OD_{600}$ ~1. Then, the whole cultures were added to 450 mL 2×YT medium + Km in 2-L baffled Erlenmeyer flasks and grown at 37°C and 250 rpm to $OD_{600}$ = 0.6-0.8, after which 1 mM IPTG was added and the cultures incubated for a further 24 h at 16°C and 150 rpm. Cells were then harvested by centrifugation and pellets frozen until protein extraction. This was performed by resuspending cells in 20 mL of His A buffer and disrupting in an Avestin Emulsiflex C5 French press (ATA Scientific Instruments).

Following centrifugation and filtration of cell extracts through 0.2-μm membranes, purification of fluorinases was carried out using 1-2 mL of HisPur Ni-NTA Resin (Thermo Scientific) in 10-mL Pierce™ disposable columns. Non-bound proteins were washed with 20 mL His A buffer before elution with 4 mL His B buffer (300 mM NaCl and 500 mM imidazole in 10 mM HEPES, pH = 7.5). The buffer of the eluted fraction was exchanged to 10 mM HEPES, pH = 7.5, using Amicon-Ultra 10,000 Da molecular-weight cut-off centrifugal units (Millipore). Protein concentration was assessed in a Nanodrop, assuming 1 $AU_{280}$ (absorbance unit measured at 280 nm) = 1 mg protein/mL. For long-term storage, 1 mM dithiothreitol (DTT) was added to the purified proteins, which were aliquoted, flash-frozen and stored at –80°C until further use.

**Fluorinase activity assays.** In these experiments, 96-well activity assays were carried out in 150 μL reactions containing 50 mM HEPES (pH = 7.8), 75 mM KF, ~1 μM fluorinase (considering the theoretical molecular mass of the protein) and 10, 25, 50, 100, 250 or 500 μM SAM (New England Biolabs). Then, the 96-well PCR plates were sealed and incubated for 1 h at 37°C followed by 5 min at 95°C, and finally cooled to 10°C in a thermocycler (Bio-Rad). The plates were then centrifuged for 30 min at 2,500×*g*, and 120-μL supernatant aliquots were transferred to round-bottom 96-well plates (Thermo Scientific), which were sealed with a silicone lid for HPLC analysis. Transhalogenation reactions were carried out in the same way, but using increasing concentrations of 5'-chloro-5'-deoxyadenosine (5'-ClDA) in the presence of 1 mM L-Met.

Steady-state kinetics were performed in the same conditions with slight modifications. Triplicate reactions (600 μL final volume) were prepared in 1.5-mL tubes and incubated for 5 min at 37°C before addition of SAM (1,5625 to 800 μM), after which 90-μL aliquots were removed at different time-points (2, 5, 10, 15, 20 and 30 min), boiled for 5 min at 95°C and placed on ice. Samples were centrifuged at maximum speed for 10 min at 4°C before being transferred to 96-well plates for 5'-FDA quantitation by HPLC analysis. Product formation rates for the different substrate concentrations were calculated by least squares linear regression with at least three time-points and fitted to the Michaelis-Menten equation using the Origin software (OriginLab Corporation).

5'-Fluoro-5'-deoxyadenosine (5'-FDA) was quantified following absorbance at 230 nm using a Zorbax C18 column (3.5 μm×4.6×100 mm, Agilent) connected to a HPLC system (Dionex Ultimate 3000, ThermoFisher) with the following gradient (at an 1 mL/min flow rate): 5-12% solvent B in 1.5 min, 12% solvent B for 1 min, 12–30% solvent B in 2 min and 30-70% B in 1.5 min [solvent A: 0.05% (v/v) acetic acid in water; solvent B: acetonitrile]. A six-point calibration curve ($R^2$ >0.99) was prepared with 0.5-50 μM 5'-FDA, which was chemically synthesized as previously described (Calero *et al.*, 2020).

***In vitro* and *in vivo* biofluorination assays in *Pseudomonas putida*.** Synthetic *flA*$^{PtaU1}$, *flA*$^{SAJ15}$, *flA*$^{Sxin}$ and *flA*$^{MA37}$ genes, codon-optimized for proper expression in Gram-negative hosts, were amplified by PCR and cloned into a pSEVA231 backbone (Silva-Rocha *et al.*, 2013) under control of the T7 promoter [plasmid termed pFB·1 in Calero *et al.* (2020)] by digestion with NdeI and SacI and ligation with T4 DNA ligase (**Tables S4** and **S5**). The resulting plasmids, encoding *N*-terminal His-tag fusions, were transformed into an engineered *P. putida* strain harboring a chromosomal copy of the T7 RNA polymerase gene under control of a fluoride-responsive riboswitch, *P. putida att*Tn*7*[FRSv1→*T7RNAP*] (Calero *et al.*, 2020). Strains carrying the different plasmids were grown overnight in de Bont medium (Hartmans *et al.*, 1989) with 5 g/L glucose and 50 mg/L Km at 30°C with shaking, and then diluted to an $OD_{600}$ = 0.1 in 50 mL of the same medium placed in 250-mL Erlenmeyer flasks. Cells were grown at 30°C with shaking of 180 rpm to an $OD_{600}$ of 0.4-0.6, at which point 15 mM NaF was added to the cultures to induce expression of the fluorinase genes. After 20 h of incubation, 2-mL samples were removed for metabolite extraction (*in vivo* assay), while the remaining cultures were centrifuged at 4,500×*g* for 10 min at 4°C to harvest cells for the *in vitro* assays.

For metabolite extraction for the *in vivo* assays, cells were centrifuged at 5,000×*g* for 10 min and the supernatants were discarded. Cells were washed once with 50 mM phosphate buffer (pH = 7.8) and incubated with 2 mL of extraction solution (40:40:20 acetonitrile:methanol:water) at −20°C for 5 min and vortexed. Cells were subsequently centrifuged at 19,000×*g* for 1 min and the supernatant was transferred to a new tube for evaporation in a SpeedVac concentrator (Thermo-Fisher Scientific) until complete dryness. Pellets were resuspended in 100 μL of Milli-Q water, centrifuged at 19,000×*g* for 1 min and the supernatant was analyzed for the presence of 5'-FDA by LC-MS as previously described (Calero *et al.*, 2020; Kittilä *et al.*, 2022).

For the *in vitro* biofluorination assays, frozen (−20°C) cell pellets were thawed, washed with 50 mM Tris·HCl buffer, pH = 7.8, and resuspended in a final volume of 3 mL of the same buffer. Cells were then disrupted with glass beads in a Precellys 24 homogenizer with two 20 s cycles at 6,000×*g*. The suspension was centrifuged at 17,000×*g* for 2 min at 4°C and the supernatant transferred to a new tube. A reaction mixture of a total volume of 100 μL was prepared with 50 μL cell-free extract, 200 μM SAM and 5 mM NaF in 50 mM Tris·HCl buffer, pH = 7.8. These reactions were incubated for 20 h at 30°C in a thermocycler, after which they were stopped with a 5-min incubation at 95°C and centrifugation for 10 min at 17,000×*g* at 4°C. Supernatants were analyzed for the presence of 5'-FDA by LC-MS as indicated above.

**Table S1.** Amino acid sequences of putative fluorinases.

| Accession no. | Organism | Abbreviation | Sequence |
|---|---|---|---|
| CDH39444.1 | *Streptomyces* sp. MA37 | FIA$^{\mathrm{MA37}}$ | MAANGSQRPIIAFMSDLGTTDDSVAQCKGLMHSICPGVTVVDVCHSMTP<br>WDVEEGARYIVDLPRFFPEGTVFATTTYPATGTTTRSVAVRIRQAAKGGA<br>RGQWAGSGDGFERADGSYIYIAPNNGLLTTVLEEHGYIEAYEVTSTKVIPA<br>NPEPTFYSREMVAIPSAHLAAGFPLAEVGRRLDDSEIVRFHRPAVEISGEA<br>LSGVVTAIDHPFGNIWTNIHRTDLEKAGIGQGKHLKIILDDVLPFEAPLTPTF<br>ADAGAIGNIAFYLNSRGYLSLARNAASLAYPYNLKAGLKVRVEAR |
| 1RQP_A | *Streptomyces cattleya* | FIA$^{\mathit{Scat}}$ | MAANSTRRPIIAFMSDLGTTDDSVAQCKGLMYSICPDVTVVDVCHSMTP<br>WDVEEGARYIVDLPRFFPEGTVFATTTYPATGTTTRSVAVRIKQAAKGGA<br>RGQWAGSGAGFERAEGSYIYIAPNNGLLTTVLEEHGYLEAYEVTSPKVIP<br>EQPEPTFYSREMVAIPSAHLAAGFPLSEVGRPLEDHEIVRFNRPAVEQDG<br>EALVGVVSAIDHPFGNVWTNIHRTDLEKAGIGYGARLRLTLDGVLPFEAPL<br>TPTFADAGEIGNIAIYLNSRGYLSIARNAASLAYPYHLKEGMSARVEAR |
| WP_019711456.1 | *Streptomyces xinghaiensis* | FIA$^{\mathit{Sxin}}$ | MSADPTQRPIIGFMSDLGTTDDSVAQCKGLMHSICPGVTVIDVCHSMTPW<br>DVEEGARYIVDLPRFFPEGTVFATTTYPATGTETRSVAVRIKQAAKGGAR<br>GQWAGSAGGFERAEGSYIYVAPNNGLLTTVLEEHGYIEAYEVSSTKVIPE<br>RPEPTFYSREMVAIPAAHLAAGFPLSEVGRPLEDSEIVRYQPPQVEISGDT<br>LTGVVSAIDHPFGNVWTNIHRTHLEKAGIGYGKRIKIILDDVLPFEQTLVPT<br>FADAGEIGGVAAYLNSRGYLSLARNAASLAYPFNLKAGLKVRVETN |
| WP_144383880.1 | *Streptomyces* sp. SAJ15 | FIA$^{\mathrm{SAJ15}}$ | **MTSNGAHRPIIAF**MSDLGTTDDSVAQCKGLMLSICPDVTIVDVCHSMTPF<br>DVEEGSRYIVDLPRFFPEGTVFATTTYPATGTETRSVAVRIRQAAQGGAR<br>GQWAGSGAGFERQEGSYIYIAPNNGLLTPVLEEHGYIEAYEVSSTEVIPE<br>RPEPTFYSREMVAIPSAHLAAGFPLDQVGRPLKDSEIVRFSRPAVETAGT<br>ELTGVVSAIDHPFGNIWTNIHRTDLEKAGIGYGRQIRITLDDVLPFELTLVPT<br>FADAGEVGNVVAYLNSRGYLSLARNAASLAYPYNLKAGLSVRVDAH |
| WP_041832040.1 | *Actinoplanes* sp. N902-109 | FIA$^{\mathrm{N902}}$ | MPANGNPIIAFMSDLGTTDDSVAQCKGLMLSICPGVTIVDVNHSMTPWDV<br>EEGARYIVDLPRFFPEGTVFATTTYPATGTATRSVALRIKQAAQGGARGQ<br>WAGSGAGFERAEGSYIYIAPNNGLLTTVIEEHGYIEAYEVSNTKVIPAEPE<br>PTFYSREMVAIPSAHLAAGFPLNEVGRALSDDEIVRFAKPKPSTVSGGVL<br>SGVITNIDHPFGNLWTNIHRTDLEKAGIGYQTQLRLLLDGVLTFDLPLVPTF<br>ADAGQIGDPVIYINSRGYLALARNAAPLAYPYNLKAGLTVTVTKA |
| SDK90789.1 | *Actinopolyspora mzabensis* | FIA$^{\mathit{Amza}}$ | MSDSYSRPIIAFMSDLGTTDDSVAQCKGLMMSICQDVTVVDVCHSMEPW<br>NVEEGARYIVDLPRFFPEGTVFATTTYPATGTTARSVAVRIKYPAKGGAR<br>GQWAGSGEGFERSEGSYIYIAPNNGLLTTVLQEHGYTEAYEVSSTDVVP<br>ARPEPTFYSREMVAIPSAHLAAGYPLEKVGRKLQDSEIVRFTPPQATVSP<br>EGDLSGVVTAIDHPFGNIWTSIHRDNLESAGVGYGTNLKIVLDDVFPFELP<br>LSPTFADAGEVGDPVVYVNSRGYLSLARNAASLAYPYNLKEGMSVRVTR<br>S |
| WP_145935427.1 | *Amycolatopsis bartoniae* | FIA$_{\mathrm{Abar}}$ | **MATTKRPVLAF**MSDLGITDDSVAQCKGLMLSICPDVTIVDVCHTMTPWDV<br>EEGARYIVDLPRFFPEGTVFATTTYPATGTGTRSVALRIKQAAKGGARGQ<br>WAGSGAGFERAEGSYIYIAPNNGLLTSVIEEHGYLEAYEVSSTEVIPEQPE<br>PTFYSREMVAVPSAHLAAGFPLEKVGRKLADDEIVRFDRPKPVQDEDGD<br>LVGVVTTIDHPFGNVWTNIHREDLEKAGVGYGTELTITLDEVLPFRLPLSP<br>TFADAGPIGSPVAYLSSRGYLALARNAASLAYPYNIQAGIPVRVHVA |
| WP_103354124.1 | *Amycolatopsis* sp. CA-128772 | FIA$^{\mathrm{CA12}}$ | MAKPSRPIIAFMSDLGITDDSVAQCKGLMLSVCPDVTIVDVCHTMKPWDV<br>EEGARYIVDLPRLFPEGTVFATTTYPATGTTTRSVALRIKQAAKGGARGQ<br>WAGSGAGFERAEGSYIYIAPNNGLLTSVIEEHGYVEAYEVSSTEVIPEQPE<br>PTFYSREMVALPSAHLAAGFPLEKVGRPLADDEIVRFERAKPAQNDDGEL<br>VGVVTAIDHPFGNVWTNIHREDLEKLGAGYGTRLRITLDEVLPFDLPLSPT<br>FADAGPIGTPVAYLSSRGYLALARNAASLAYPYNLNAGISVRVVAA |
| WP_149854061.1 | *Goodfellowiella* sp. AN110305 | FIA$^{\mathrm{AN11}}$ | MASSRGNRPIIAFMSDLGITDDSVAQCKGLMLSVCPDVTIVDVCHTMKPW<br>DVEEGARYIVDLPRFFPEGTVFATTTYPATGTTARSVAIRIKQAAKGGARG<br>QWAGSGAGFERAEGSYIYIAPNNGLLTSVIEEHGYLEAYEVSSTEVIPELP<br>EPTFYSREMVALPSGHLAAGFPLNKVGRPLADDEIVRFTRPQPSVETDGD<br>LVGVVTNIDHPFGNIWTNIHRTDLERAGVGYGTKLKVVLDEVLPFELPLSP<br>TFADAGPVGTPVAYLSSRGYLALARNAASLAYPYNLEAGIPVRVKVG |
| WP_029901962.1 | *Nocardia brasiliensis* IFM 10847 | FIA$^{\mathit{Nbra2}}$ | **MTTANGRRPIIAF**MSDLGITDDSVAQCKGLMLSVCPDVTIVDICHTMQPW<br>DVEEGARYIVDLPRLFPEGTVFATTTYPATGTTARSVALRIAHASKGGAR<br>GQWAGSGAGFERKEGSYIYIAPNNGLLTTVIKEHGYLEAYEVSSPEVIPEQ<br>PEPTFYSREMVALPSAHLAAGFPLEKVGRRLADDEIVRFERKDPELVADN<br>ELLGYVTNIDHPFGNVWTNIHRTDLEKLGVGYGTELRITLDGVLPFELPLS<br>PTFADAGEVGAAVAYLSSRGYLALARNAASLAYPYNLNAGISVQVKVD |

| | | | |
|---|---|---|---|
| SUB47361.1 | *Nocardia brasiliensis* NCTC 11294 | Fla$^{Nbra3}$ | MTTANGRRPIIAFMSDLGITDDSVAQCKGLMLSVCPDVTIVDICHTMQPW<br>DVEEGARYIVDLPRLFPEGTVFATTTYPATGTTARSVALRIAHASKGGAR<br>GQWAGSGAGFERKEGSYIYIAPNNGLLTTVIKEHGYLEAYEVSSPEVIPEQ<br>PEPTFYSREMVALPSAHLAAGFPLEKVGRRLADDEIVRFERKDPELVADT<br>DLVGYVTNIDHPFGNVWTNIHRTDLEKLGVGYGTKLRITLDGVLPFELPLS<br>PTFADAGEVGAAVAYLSSRGYLALARNAASLAYPYNLKAGISVQVKVG |
| AHK61118.1 | *Nocardia brasiliensis* ATCC 700358 | FlA$^{Nbra1}$ | MTTTNGRRPIIAFMSDLGITDDSVAQCKGLMLSVCPDVTIVDICHTMQPW<br>DVEEGARYIVDLPRLFPEGTVFATTTYPATGTTARSVALRIAHASKGGAR<br>GQWAGSGAGFERKEGSYIYIAPNNGLLTTVIKEHGYLEAYEVSSPEVIPEQ<br>PEPTFYSREMVALPSAHLAAGFPLEKVGRRLADDEIVRFERKDPELVADH<br>DLVGYVTNIDHPFGNVWTNIHRTDLEKLGVGYGTKLRITLDGVLPFELPLS<br>PTFADAGEIGAAVAYLSSRGYLALARNAASLAYPYNLKAGISVQVKVG |
| TMD14026.1 | *Chloroflexi bacterium* | FlA$^{Cbac}$ | **MQEGQATSTPRQRPVIAF**MSDLGTFDDSVGICKGLMLSVCPDVVIVDICH<br>AMTPFDIEEGARLIVDLPRFFPEGTVFATTTYPATGTSTRSVALRIRQAAV<br>GGARGQWAGAGEGIQRAEGAYIYIAPNNGLLTSVIEEHGYLEAYEVTSTS<br>VIPARPEPTFFSREMVAVPAAHLAAGFPLSQVGRPLQDSEIVRFDRRRPA<br>SLGDGGFAGVITVVDRPYGNVWTNISRKDLANQGIAYGTRLRITVDNVLP<br>FDLTLTPTFADAGEIGAPVCYVNSRGYLSLARNASNLADTYNIRRHMPVN<br>VQVVSRTDGEPHRSESSELASVKGT |
| KLU63204.1 | *Peptococcaceae bacterium* CEB3 | FlA$^{Pbac}$ | MEKVDNKTVFQRPIIAFMSDLGAFDDSVGICKGLMLSVCPEAQIIDICHSM<br>TPFDVEQGARLIVDLPRFFPEGRTVFATTTYPATGTMARSLALRIKRPAKG<br>GALGQWAGAGFGIERGVGGYIYVAPNNGLLTDVIEEHGYLEAYEITSTNVI<br>PENPEPTFFSREMVALPSAYLAAGYPLSEVGQPLGDSEIVRFKKVLPRKM<br>SESELVGVVAAIDRPFGNVWTNISRRDLDKIGVTYGSQLKVVLDNALMFE<br>LPLSQTFADAREIGAAVAYINSRGHLSLGRYAANLADRYNINRGMPIRLKVI<br>TG |
| WP_093392705.1 | *Thermodesulforhabdus norvegica* | FlA$^{Tnor}$ | MDIGPIIGFASDLGLKDDSVALCKGLMISICPEVYIVDICHTMTPFDIEEGAW<br>LALDLPRFFPEGRTIFAVTTYPATGTEARSIAVRIKKAVPGGSLEKWEGPG<br>GGMERTLEGGYIYVAPNNGLLTFVLERYGYIEAYEIISTEFIPENPEPTFFS<br>REMVAIRAACIAKKVVSQGVPLSKVGPPITEDKLARIKLSLPEKIAHNEIRG<br>KIIRIDYPYGNVWTNISFNDLKSMGINYGSRLTVVIGDILSFNVYLTRTFADA<br>GGIGDVISYINSRGYFSLGGYAANLADLCNLRRGMNVVVIKV |
| OPY51785.1 | *Methanosaeta* sp. PtaU1.Bin055 | FlA$^{PtaU1}$ | MLQNTKNEYKPRDGGGHRPIIGFMSDLGTEDDSVGICKGVMLGLCPDAVI<br>VDISHSMTPWDIDQGSRLIVDLPKFFPNWTTFATTSYRETGTSARSVAIKL<br>PSGHVYVAPNNGLLTRVIEDHGYVEAYEVTTVGAIPAEPEPTWFSRDMVA<br>YPAAAIAAGFPLENVGRPLNDSEIVRADLPRYTQAEDGIIQGIVTTIDRPFG<br>NIWTNIPRRIIENELRIAYGTNIRVVLDNLLVLEVPFMRTFGDVGLKKPMCYI<br>NSRGYFSLAYYGGNLADPYNIRRGMPVKIESITR |
| 2Q6L_A | *Salinispora tropica* CNB-440 | SalL$^{Stro}$ | MQHNLIAFLSDVGSADEAHALCKGVMYGVAPAATIVDITHDVAPFDVREG<br>ALFLADVPHSFPAHTVICATVYPETGTATHTIAVRNEKGQLLVGPNNGLLS<br>FALDASPAVECHEVLSPDVMNQPVTPTWYSKDIVAACAAHLAAGTDLAAV<br>GPRIDPKQIVRLPYASASEVEGGIRGEVVRIDRAFGNVWTNIPTHLIGSML<br>QDGERLEVKIEALSDTVLELPFCKTFGEVDEGQPLLYLNSRGRLALGLNQ<br>SNFIEKWPVVPGDSITVSPRVPDSNLGPVLG |

*N*-terminal residues added after manual curation are highlighted in bold font.

**Table S2.** NCBI accession numbers of 16S rRNA sequences used for phylogenetic analysis of organisms encoding fluorinase genes.

| Organism | Accession no. |
|---|---|
| *Streptomyces* sp. MA37 | HG428740 |
| *Streptomyces cattleya* | NC_017586.1 [2,721,506–2722810; 4,855,739–4,857,043; 3,613,701–3,615,005; 1,079,076–1,080,380; 2,144,515–2,145,819; 802,858–804,162] |
| *Streptomyces xinghaiensis* | NR_116059.1 |
| *Streptomyces* sp. SAJ15 | NISY01000001.1 [424-1,746] |
| *Actinoplanes* sp. N902-109 | CP005929.1 [183,863–185,376; 1,631,125–1,632,638; 6,234,429–6,235,942; 6,736,828–6,738,341] |
| *Actinopolyspora mzabensis* | NR_109353.1 |
| *Amycolatopsis bartoniae* | MN399889.1 |
| *Amycolatopsis* sp. CA-128772 | PPHG01000117.1 [201–1,497]; PPHG01000111.1 [717–2,013]; PPHG01000090.1 [39303–40599]; PPHG01000071.1 [231,228–232,524]; PPHG01000056.1 [158–1,454]; PPHG01000030.1 [16,431–17,727] |
| *Goodfellowiella* sp. AN110305 | KX762322.1 |
| *Nocardia brasiliensis* ATCC 700358 | CP003876.1 [1,448,256–1,449,778; 2,271,961–2,273,483] |
| *Chloroflexi* bacterium | VBIC01000091.1 [1–430] |
| *Peptococcaceae* bacterium CEB3 | DXJ01000019.1 [93–1,424] |
| *Thermodesulforhabdus norvegica* | NR_025970.1 |
| *Methanosaeta* sp. PtaU1.Bin055 | AJ133791.1 |
| *Salinispora tropica* CNB-440 | NR_074502.1 |

The organisms used in this analysis are as shown in **Figure S1**. Numbers between brackets indicate nucleotide positions

**Table S3.** NCBI accession numbers for the nucleotide sequences harboring *fl* gene clusters.

| Organism | Accession no. |
|---|---|
| *Streptomyces* sp. MA37 | HG428738.2 |
| *Streptomyces cattleya* | NC_016111; NC_016113 |
| *Streptomyces xinghaiensis* | NZ_CP023202 |
| *Streptomyces* sp. SAJ15 | NZ_NISY01000005; NISY01000015 |
| *Actinoplanes* sp. N902-109 | NC_021191 |
| *Actinopolyspora mzabensis* | FNFM01000012; FNFM01000017 |
| *Amycolatopsis bartoniae* | NZ_JACHWH010000014 |
| *Amycolatopsis* sp. CA-128772 | NZ_PPHG01000085 |
| *Goodfellowiella* sp. AN110305 | NZ_VUOB01000072 |
| *Nocardia brasiliensis* IFM 10847 | NZ_BAUA01000190 |
| *Nocardia brasiliensis* NCTC 11294 | UGSN01000003 |
| *Nocardia brasiliensis* ATCC 700358 | NC_018681 |
| *Chloroflexi* bacterium | VBIC01000085 |
| *Peptococcaceae* bacterium CEB3 | LDXJ01000028 |
| *Thermodesulforhabdus norvegica* | NZ_FOUU01000001 |
| *Methanosaeta* sp. PtaU1.Bin055 | MVRM01000143 |
| *Salinispora tropica* CNB-440 | NC_009380 |

**Table S4.** Nucleotide sequences of synthetic genes encoding fluorinases.

| Description | Sequence (5'→3') |
|---|---|
| *flA* from *Streptomyces* sp. MA37 (*flA*$^{MA37}$) codon optimized for expression in *P. putida* | ATGGCCGCCAACGGCAGCCAGCGCCCGATCATCGCGTTCATGTCCGACCTGGGCACCACCGACGACAGCGTGGCGCAG<br>TGCAAGGGCCTGATGCACAGCATCTGCCCCGGTGTGACCGTGGTGGATGTGTGCCACAGCATGACCCCGTGGGACGTC<br>GAGGAGGGCGCCCGTTACATCGTGGACCTGCCGCGTTTCTTCCCGGAGGGCACCGTCTTCGCCACCACCACCTATCCC<br>GCCACCGGCACCACCACCCGCAGCGTGGCCGTGCGTATCCGCCAGGCCGCCAAAGGCGGCGCCCGTGGCCAGTGGGCC<br>GGCAGCGGCGACGGCTTCGAACGTGCCGACGGCAGCTACATCTACATCGCCCCGAACAACGGCCTGCTGACCACCGTG<br>CTGGAGGAACACGGCTATATCGAGGCCTACGAGGTGACCAGCACCAAGGTGATCCCGGCGAACCCGGAGCCGACCTTC<br>TACAGCCGCGAAATGGTGGCCATCCCGTCCGCCCACCTGGCCGCCGGCTTCCCGCTGGCCGAGGTGGGCCGTCGTCTG<br>GATGACAGCGAGATCGTCCGTTTCCACCGCCCCGCCGTCGAGATCTCCGGCGAAGCCCTGAGCGGCGTGGTGACCGCC<br>ATCGACCACCCGTTCGGCAACATCTGGACCAACATCCACCGTACCGACCTGGAAAAGGCCGGCATCGGCCAGGGCAAA<br>CACCTGAAGATCATCCTGGACGACGTGCTGCCGTTCGAAGCCCCGCTGACCCCCACCTTCGCCGACGCCGGCGCCATC<br>GGCAACATCGCCTTCTACCTGAACAGCCGCGGCTATCTGAGCCTGGCCCGCAACGCCGCCAGCCTGGCCTATCCGTAC<br>AACCTGAAAGCCGGCCTGAAGGTGCGCGTGGAGGCCCGTTGA |
| *flA* from *Streptomyces cattleya* (*flA*$^{Scat}$) codon optimized for expression in *P. putida* | ATGGCCGCCAACAGCACCCGCCGCCCGATCATCGCCTTCATGAGCGACCTGGGCACCACCGACGACAGCGTGGCCCAG<br>TGCAAGGGCCTGATGTACAGCATCTGCCCGGACGTGACCGTGGTGGATGTGTGCCACTCCATGACCCCGTGGGACGTG<br>GAGGAGGGCGCCCGTTACATCGTGGACCTGCCGCGTTTCTTCCCGGAGGGCACCGTGTTCGCCACCACCACCTATCCC<br>GCGACCGGCACCACCACCCGCAGCGTGGCCGTGCGTATCAAACAGGCCGCCAAAGGCGGCGCCCGCGGCCAGTGGGCC<br>GGCAGCGGCGCCGGCTTCGAGCGTGCCGAGGGCAGCTACATCTACATCGCCCCCAACAACGGCCTGCTGACCACCGTG<br>CTGGAGGAACATGGCTACCTGGAGGCCTACGAAGTGACCAGCCCGAAGGTGATCCCGGAGCAGCCGGAACCGACCTTC<br>TACAGCCGCGAGATGGTGGCCATCCCGTCCGCCCACCTGGCCGCCGGCTTCCCGCTGAGCGAAGTGGGCCGTCCGCTG<br>GAAGACCACGAAATCGTCCGCTTCAACCGCCCGGCCGTGGAGCAGGACGGCGAAGCCCTGGTGGGCGTCGTGAGCGCC<br>ATCGATCACCCGTTCGGCAACGTGTGGACCAACATCCACCGCACCGACCTGGAGAAGGCCGGCATCGGCTACGGCGCC<br>CGTCTGCGTCTGACCCTGGACGGCGTGCTGCCGTTCGAAGCCCCGCTGACCCCGACCTTCGCCGACGCCGGCGAAATC<br>GGCAACATCGCCATCTACCTGAACAGCCGCGGCTACCTGTCCATCGCCCGCAACGCCGCCTCGCTGGCCTATCCGTAC<br>CACCTGAAGGAGGGCATGAGCGCCCGCGTGGAGGCCCGTTGA |
| *flA* from *Streptomyces xinghaiensis* (*flA*$^{Sxin}$) codon optimized for expression in *P. putida* | ATGTCCGCCGATCCAACACAACGTCCAATTATAGGTTTCATGAGCGACTTGGGCACCACCGACGACAGTGTAGCCCAG<br>TGCAAAGGCCTGATGCACTCGATCTGCCCTGGGGTAACAGTTATCGATGTGTGCCACAGCATGACCCCGTGGGACGTG<br>GAGGAGGGTGCCCGCTACATCGTGGACCTGCCGCGCTTCTTCCCAGAGGGCACCGTGTTCGCCACCACAACCTATCCG<br>GCCACTGGGACCGAAACCCGTTCGGTGGCGGTGCGTATCAAGCAGGCCGCCAAGGGCGGCGCTCGCGGGCAGTGGGCC<br>GGCAGCGCCGGTGGCTTCGAAAGGGCGGAAGGCAGCTACATCTACGTGGCCCCGAACAACGGCCTGCTGACCACCGTA<br>CTCGAAGAGCATGGCTATATCGAGGCCTACGAGGTGAGTTCCACCAAGGTCATCCCGGAGCGCCCGGAACCCACGTTC<br>TACTCGCGAGAAATGGTCGCCATTCCGGCAGCGCATTTAGCCGCGGGGTTCCCGCTGTCGGAAGTGGGTCGCCCTTTG<br>GAAGACAGCGAGATCGTCCGTTACCAGCCGCCGCAGGTGGAAATCAGCGGCGACACCCTCACCGGCGTGGTCTCAGCC<br>ATCGACCACCCGTTCGGCAACGTGTGGACCAACATCCACCGCACCCACCTGGAAAAAGCCGGCATCGGCTACGGCAAG<br>CGCATCAAGATCATCCTGGACGACGTTCTGCCCTTCGAGCAGACCCTGGTGCCGACCTTCGCCGACGCCGGCGAGATT<br>GGCGGGGTTGCGGCCTATCTCAACAGCCGCGGCTACCTGAGCCTGGCCCGCAACGCCGCTTCGCTGGCTTACCCGTTC<br>AACCTGAAGGCCGGGCTCAAGGTGCGCGTTGAAACCAACTGA |
| *flA* from *Streptomyces* sp. SAJ15 (*flA*$^{SAJ15}$) codon optimized for expression in *E. coli* | ATGACGTCCAACGGTGCCCATCGCCCCATCATCGCGTTTATGTCCGATTTAGGGACCACAGATGATTCCGTTGCTCAG<br>TGTAAAGGCCTTATGTTATCGATTTGTCCCGATGTAACCATTGTCGATGTCTGTCACAGTATGACCCCGTTCGATGTA<br>GAGGAAGGTTCGCGCTACATTGTTGACCTGCCGCGTTTTTTTCCCGAGGGCACCGTTTTTGCTACTACGACATATCCT<br>GCTACCGGTACTGAAACCCGTAGTGTTGCGGTTCGTATTCGTCAAGCAGCACAAGGTGGAGCGCGCGGACAGTGGGCT<br>GGTTCCGGAGCCGGGTTCGAGCGCCAAGAAGGTAGCTACATCTATATCGCGCCAAACAACGGGCTTCTTACGCCGGTC<br>TTGGAGGAACACGGATACATCGAAGCTTACGAGGTGTCAAGCACCGAAGTAATCCCTGAACGTCCTGAACCAACCTTC<br>TACTCACGCGAGATGGTCGCGATTCCGTCCGCTCATTTGGCCGCCGGATTTCCACTTGACCAGGTGGGTCGTCCGCTG<br>AAGGACTCCGAAATCGTCCGTTTCAGCCGCCCGGCGGTCGAAACAGCGGGCACGGAATTGACAGGGGTGGTGTCAGCT<br>ATCGATCACCCATTTGGTAATATCTGGACTAACATCCATCGCACTGACTTAGAGAAGGCTGGGATCGGCTATGGCCGC<br>CAGATTCGCATCACTCTTGACGACGTATTACCGTTTGAATTGACTCTTGTCCCTACATTTGCTGACGCTGGCGAGGTT<br>GGAAATGTAGTGGCTTATTTGAATTCGCGCGGATACCTGAGTTTAGCACGCAACGCTGCGTCACTTGCCTATCCGTAC<br>AACCTGAAAGCCGGACTGTCGGTGCGCGTGGACGCTCATTAA |
| *flA* from *Actinoplanes* sp. N902-109 (*flA*$^{N902}$) codon optimized for expression in *E. coli* | ATGCCCGCAAACGGAAACCCTATCATCGCATTTATGTCTGACCTGGGTACAACTGATGATTCGGTCGCGCAGTGTAAA<br>GGGTTAATGCTGTCAATTTGTCCCGGTGTTACCATCGTGGACGTAAACCATTCTATGACTCCATGGGACGTGGAGGAG<br>GGAGCCCGCTATATCGTTGACTTACCTCGTTTTTTCCCGGAGGGTACAGTTTTCGCCACGACGACTTATCCCGCCACT<br>GGTACGGCAACCCGTAGCGTTGCTTTGCGTATCAAACAAGCAGCTCAAGGCGGGGCTCGCGGACAGTGGGCTGGTTCC<br>GGAGCCGGCTTCGAACGCGCAGAAGGCTCATACATTTATATTGCCCCTAACAATGGGCTGTTAACTACGGTCATCGAA<br>GAACACGGCTATATTGAGGCTTACGAAGTATCTAATACGAAAGTTATTCCAGCAGAGCCGGAGCCAACATTTTACAGC<br>CGCGAAATGGTGGCTATTCCTTCTGCCCACCTGGCAGCGGGTTTTCCACTTAATGAAGTTGGACGCGCACTTAGCGAC<br>GACGAAATCGTCCGCTTCGCAAAACCAAAGCCGTCAACAGTTAGTGGAGGCGTCTTATCCGGTGTCATCACCAATATT<br>GACCATCCATTCGGTAATCTGTGGACTAATATTCACCGTACGGATCTGGAGAAGGCAGGTATCGGCTACCAAACGCAA<br>TTACGCTTGCTGCTTGACGGGGTCCTGACCTTTGATCTGCCGTTAGTACCAACTTTCGCCGACGCGGGACAGATCGGG<br>GATCCGGTCATCTACATCAACTCACGCGGGTATTTGGCGTTGGCTCGTAACGCGGCACCATTGGCCTATCCCTACAAC<br>TTAAAGGCGGGTTTAACGGTGACAGTGACTAAA |

| | |
|---|---|
| *flA* from *Actinopolyspora mzabensis* (*flA*$^{Amza}$) codon optimized for expression in *E. coli* | ATGTCGGACAGTTATTCGCGGCCGATCATCGCTTTCATGAGCGACCTGGGCACTACTGATGATTCGGTCGCGCAGTGT<br>AAAGGCCTGATGATGAGCATATGCCAGGATGTGACCGTAGTGGACGTGTGCCACTCCATGGAGCCCTGGAATGTGGAG<br>GAGGGAGCTCGCTACATCGTTGACCTGCCCCGGTTCTTCCCCGAAGGCACCGTCTTCGCCACCACCACTTATCCGGCC<br>ACCGGCACTACGGCGCGATCGGTGGCGGTACGGATCAAATATCCGGCCAAAGGCGGTGCTCGTGGTCAGTGGGCGGGG<br>TCCGGCGAGGGATTCGAACGCAGCGAGGGCTCCTATATCTACATCGCCCCCAACAACGGTCTTCTCACGACTGTGCTC<br>CAGGAGCACGGTTATACGGAGGCATATGAGGTGAGCTCGACGGATGTCGTTCCCGCCAGGCCTGAACCGACTTTCTAC<br>AGTCGGGAAATGGTTGCCATCCCATCAGCGCATCTTGCGGCGGGGTATCCGTTGGAAAAGGTGGGGCGCAAGCTGCAG<br>GACTCCGAGATAGTCCGGTTCACGCCACCACAGGCCACGGTCTCACCAGAGGGCGATCTAAGCGGTGTCGTAACAGCA<br>ATAGACCATCCCTTCGGCAATATCTGGACCAGCATTCACCGGGACAACCTCGAATCTGCCGGAGTCGGTTACGGCACG<br>AACCTGAAAATCGTTCTGGACGACGTTTTCCCCTTCGAATTGCCGTTGAGCCCCACATTCGCCGATGCGGGTGAAGTT<br>GGTGATCCGGTCGTTTATGTCAACAGTCGTGGTTATCTGTCCTTGGCCCGCAACGCGGCAAGTCTGGCCTACCCGTAC<br>AACCTGAAAGAAGGAATGTCAGTCAGAGTGACCCGGTCGTAG |
| *flA* from *Amycolatopsis bartoniae* (*flA*$^{Abar}$) codon optimized for expression in *E. coli* | ATGGCTACTACGAAGCGCCCAGTCCTTGCTTTCATGAGTGATCTGGGCATCACGGACGATAGTGTTGCACAGTGTAAA<br>GGATTAATGCTGTCGATCTGTCCTGACGTTACAATCGTCGATGTGTGCCACACCATGACGCCGTGGGACGTGGAGGAG<br>GGCGCACGTTACATTGTCGACTTGCCACGCTTTTTCCCAGAGGGGACCGTCTTCGCCACCACAACTTATCCGGCTACG<br>GGTACGGGTACACGTTCCGTGGCCTTGCGCATTAAGCAGGCCGCAAAGGGAGGAGCTCGTGGACAATGGGCTGGCTCT<br>GGGGCTGGCTTCGAACGTGCAGAGGGCAGTTATATCTACATCGCTCCGAACAATGGGTTGTTGACAAGTGTAATCGAG<br>GAACATGGGTACCTGGAAGCATACGAAGTGTCGAGCACAGAGGTTATTCCTGAACAACCCGAACCTACCTTTTACTCA<br>CGCGAGATGGTGGCCGTTCCAAGTGCCCACCTTGCCGCAGGGTTTCCATTAGAGAAAGTAGGCCGCAAATTAGCAGAT<br>GATGAGATTGTACGCTTCGACCGTCCTAAACCAGTACAAGACGAGGATGGTGACTTAGTAGGAGTAGTAACTACAATT<br>GATCACCCTTTTGGCAATGTTTGGACCAATATTCATCGTGAAGATCTTGAAAAAGCTGGTGTCGGCTATGGGACTGAG<br>CTTACTATCACCCTGGACGAAGTCCTGCCGTTTCGCCTTCCCTTGTCGCCCACTTTCGCGGACGCGGGACCAATCGGG<br>TCTCCTGTAGCCTATTTAAGTTCACGCGGGTACCTTGCTTTAGCCCGCAACGCCGCAAGCCTGGCGTACCCTTATAAT<br>ATCCAGGCTGGGATTCCTGTCCGCGTGCACGTCGCTTAA |
| *flA* from *Amycolatopsis* sp. CA-128772 (*flA*$^{CA12}$) codon optimized for expression in *E. coli* | ATGGCAAAGCCCTCGCGCCCTATCATTGCCTTTATGAGCGACCTGGGGATCACCGATGATTCGGTCGCGCAGTGTAAG<br>GGGCTCATGTTGAGCGTCTGCCCAGATGTCACCATCGTGGATGTCTGTCACACGATGAAGCCGTGGGACGTGGAGGAA<br>GGCGCTCGGTACATCGTCGATCTGCCCCGGCTCTTTCCGGAGGGGACGGTGTTCGCAACCACGACGTATCCAGCAACG<br>GGTACCACCACCCGTTCCGTGGCCCTGCGCATTAAACAAGCGGCGAAGGGTGGGGCTCGTGGTCAATGGGCAGGTTCC<br>GGGGCCGGCTTTGAGCGCGCGGAAGGTTCGTATATCTACATCGCGCCTAACAACGGGTTGCTCACCTCGGTCATCGAA<br>GAACACGGCTATGTCGAAGCTTATGAGGTGAGCTCCACCGAAGTGATCCCTGAGCAGCCCGAACCTACCTTTTATAGC<br>CGGGAAATGGTCGCGTTGCCAAGCGCGCACTTGGCAGCAGGCTTCCCACTCGAGAAAGTGGGTCGCCCGTTGGCCGAT<br>GACGAAATCGTGCGGTTCGAGCGGGCTAAACCAGCACAAAATGACGATGGCGAGTTGGTGGGTGTCGTGACCGCTATC<br>GATCACCCTTTTGGGAATGTGTGGACCAATATTCACCGCGAGGACCTCGAGAAGCTCGGTGCGGGCTACGGTACCCGC<br>CTCCGCATCACCTTGGACGAGGTCTTGCCGTTTGACCTGCCGCTGTCCCCAACCTTTGCAGATGCAGGGCCAATTGGG<br>ACCCCAGTCGCGTATCTGTCGTCGCGTGGTTATTTGGCACTCGCGCGTAACGCGGCTAGCTTGGCTTACCCTTATAAT<br>TTGAACGCGGGTATCTCGGTCCGGGTCGTCGCCGCCTGA |
| *flA* from *Goodfellowiella* sp. AN110305 (*flA*$^{AN11}$) codon optimized for expression in *E. coli* | ATGGCATCTTCGCGTGGTAATCGCCCGATTATCGCTTTCATGTCTGACTTGGGCATTACCGATGATTCTGTGGCACAA<br>TGCAAGGGGCTGATGTTGTCCGTTTGTCCCGACGTTACAATCGTAGACGTCTGCCATACGATGAAGCCGTGGGATGTA<br>GAGGAAGGCGCGCGTTACATCGTAGACCTGCCTCGTTTTTTTCCCGAAGGAACAGTGTTTGCGACGACCACTTATCCT<br>GCAACGGGGACTACTGCGCGTTCTGTAGCAATTCGTATTAAACAAGCGGCGAAGGGTGGGGCTCGTGGTCAATGGGCC<br>GGATCGGGAGCCGGTTTCGAGCGCGCTGAAGGGTCGTACATTTATATCGCACCTAACAACGGACTGCTGACATCAGTT<br>ATTGAGGAACACGGCTATTTAGAAGCCTACGAAGTTTCGAGTACAGAGGTGATCCCGGAGCTGCCCGAGCCTACGTTC<br>TATAGCCGTGAGATGGTAGCGCTTCCTTCGGGCCATCTTGCAGCTGGTTTTCCATTGAACAAAGTCGGTCGTCCGTTA<br>GCAGACGATGAAATCGTGCGTTTTACACGCCCACAACCTTCAGTTGAAACCGATGGAGACCTTGTCGGGGTCGTAACC<br>AATATTGACCACCCCTTCGGAAATATTTGGACGAACATTCATCGTACTGACTTGGAGCGCGCGGGCGTAGGGTATGGG<br>ACCAAATTAAAAGTCGTATTAGACGAAGTTTTGCCTTTCGAGCTGCCTTTATCCCCCACATTCGCAGATGCGGGGCCG<br>GTAGGAACACCAGTGGCTTATCTTAGTTCACGTGGGTACTTGGCTTTGGCTCGCAACGCCGCGTCCCTGGCTTACCCG<br>TACAATCTTGAGGCTGGCATTCCAGTACGTGTAAAGGTTGGATAA |
| *flA* from *Nocardia brasiliensis* IFM 10847 (*flA*$^{Nbra2}$) codon optimized for expression in *E. coli* | ATGACCACGGCGAATGGACGTCGTCCTATTATTGCATTTATGAGCGATCTGGGGATCACTGACGATTCGGTCGCCCAG<br>TGTAAGGGTTTAATGTTGTCTGTGTGTCCTGATGTAACTATTGTTGACATTTGTCACACGATGCAGCCCTGGGACGTG<br>GAAGAGGGCGCTCGTTACATCGTTGACCTGCCTCGTCTTTTTCCCGAAGGAACTGTATTTGCGACGACCACCTATCCA<br>GCTACAGGAACTACAGCTCGTTCCGTCGCTCTGCGTATTGCTCATGCCTCGAAAGGTGGCGCACGCGGTCAATGGGCC<br>GGAAGTGGCGCAGGGTTTGAGCGCAAGGAGGGAAGCTATATCTATATTGCACCGAATAATGGCCTGCTGACTACCGTA<br>ATCAAAGAACACGGTTATCTGGAAGCGTATGAGGTGTCCTCCCCCGAAGTGATTCCCGAACAACCCGAGCCAACTTTT<br>TACAGTCGCGAGATGGTTGCCCTTCCTTCAGCGCATTTAGCCGCGGGGTTTCCTTTAGAGAAAGTAGGGCGTCGCCTT<br>GCTGATGATGAGATTGTCCGTTTTGAACGCAAAGACCCAGAACTTGTGGCAGACAACGAGTTGTTAGGCTACGTTACG<br>AACATTGACCACCCATTTGGAAATGTCTGGACTAACATCCACCGCACCGATTTGGAGAAGTTGGGGGTAGGCTATGGT<br>ACTGAATTACGCATCACGTTGGATGGTGTCTTACCATTCGAACTTCCGCTTTCACCAACCTTCGCTGACGCCGGTGAG<br>GTTGGTGCAGCCGTTGCTTATTTATCTTCACGCGGCTATTTAGCACTTGCGCGTAATGCCGCTTCACTGGCATATCCT<br>TACAACTTGAATGCAGGGATTTCAGTGCAGGTTAAGGTTGACTAA |

| | |
|---|---|
| *flA* from *Nocardia brasiliensis* NCTC 11294 (*flA*$^{Nbra3}$) codon optimized for expression in *E. coli* | ATGACTACCGCAAACGGTCGCCGCCCCATCATTGCTTTCATGTCTGATCTTGGCATCACTGACGACTCTGTGGCGCAA<br>TGTAAGGGGCTGATGTTGAGTGTTTGCCCAGACGTAACAATTGTCGATATTTGTCACACTATGCAACCGTGGGATGTC<br>GAGGAAGGTGCTCGTTATATCGTCGACTTGCCTCGTCTGTTTCCTGAAGGGACGGTCTTCGCCACAACAACTTACCCG<br>GCGACTGGAACAACAGCTCGCTCTGTGGCCCTGCGCATCGCACATGCGTCAAAAGGAGGAGCACGTGGTCAATGGGCT<br>GGTTCGGGTGCTGGTTTCGAACGTAAGGAAGGGTCTTACATTTACATTGCCCCCAACAACGGACTGTTGACCACGGTC<br>ATCAAGGAACACGGCTACCTGGAAGCATACGAAGTGAGCAGCCCGGAGGTCATTCCTGAGCAACCGGAACCTACGTTT<br>TATTCACGCGAAATGGTTGCCTTGCCGTCGGCCCATCTTGCTGCAGGTTTCCCACTTGAAAAGGTGGGTCGTCGTTTA<br>GCAGACGATGAGATCGTGCGCTTCGAACGTAAAGATCCGGAACTTGTCGCGGACACTGATTTAGTTGGCTATGTGACG<br>AATATTGACCATCCGTTTGGAAACGTCTGGACGAACATTCATCGCACCGATCTGGAAAAGTTGGGAGTGGGTTATGGA<br>ACCAAACTTCGCATCACCTTAGACGGTGTGTTACCGTTTGAGTTGCCCTTAAGCCCGACTTTTGCCGATGCCGGGGAA<br>GTCGGAGCAGCGGTGGCGTATTTGTCATCACGTGGTTATTTAGCTCTTGCTCGCAATGCCGCATCTCTGGCTTATCCG<br>TATAACCTGAAGGCGGGTATTAGTGTTCAAGTAAAAGTAGGCTAA |
| *flA* from *Nocardia brasiliensis* ATCC 700358 (*flA*$^{Nbra1}$) codon optimized for expression in *E. coli* | ATGACGACAGCTAATGGCCGTCGCCCTATCATTGCATTTATGTCTGACTTGGGGATCACTGATGATTCTGTCGCTCAA<br>TGTAAGGGGCTTATGCTTAGCGTTTGCCCTGACGTAACGATTGTGGACATCTGCCACACTATGCAGCCTTGGGATGTT<br>GAAGAAGGCGCACGCTATATTGTGGACTTACCGCGTCTGTTCCCAGAAGGTACGGTTTTCGCAACTACTACTTATCCA<br>GCTACGGGAACTACCGCACGTTCTGTTGCACTGCGTATCGCACACGCGTCAAAGGGTGGTGCCCGCGGGCAATGGGCT<br>GGGTCAGGAGCAGGGTTCGAACGTAAAGAGGGATCCTATATTTATATTGCGCCCAATAATGGACTTCTTACGACCGTC<br>ATTAAGGAACACGGTTACCTGGAAGCTTATGAGGTATCTAGTCCTGAGGTTATCCCAGAGCAACCGGAACCCACCTTT<br>TACTCTCGCGAAATGGTTGCATTACCTTCAGCTCACTTGGCTGCTGGCTTCCCCTTAGAAAAAGTTGGTCGTCGTCTT<br>GCTGACGACGAAATCGTTCGCTTTGAGCGCAAGGACCCAGAGCTGGTTGCGGACACCGACTTAGTCGGATACGTTACG<br>AATATCGACCACCCTTTCGGAAATGTTTGGACGAACATCCATCGCACGGACTTGGAAAAGTTAGGAGTGGGGTATGGA<br>ACTAAGTTACGCATCACGTTGGATGGAGTCTTACCTTTCGAACTGCCATTATCGCCGACATTTGCAGACGCGGGAGAA<br>GTGGGTGCTGCTGTCGCGTACCTTTCCTCTCGCGGCTACCTGGCATTAGCTCGCAATGCAGCATCACTTGCATATCCC<br>TACAACTTAAAAGCGGGTATCTCCGTGCAGGTTAAGGTAGGTTAA |
| *flA* from *Chloroflexi bacterium* (*flA*$^{Cbac}$) codon optimized for expression in *E. coli* | ATGCAAGAGGGGCAGGCCACATCAACTCCCCGTCAGCGCCCCGTCATTGCCTTTATGTCTGATCTGGGTACCTTTGAT<br>GACTCCGTTGGAATTTGTAAAGGACTGATGCTGAGTGTGTGCCCGGATGTGGTCATCGTGGATATCTGCCACGCCATG<br>ACCCCGTTCGACATTGAGGAGGGAGCTCGTCTGATCGTCGATTTACCTCGTTTCTTTCCCGAGGGAACTGTTTTTGCT<br>ACCACCACGTATCCAGCCACGGGTACTAGCACACGCTCTGTCGCTTTACGTATTCGTCAGGCTGCAGTAGGAGGAGCA<br>CGTGGTCAATGGGCCGGTGCAGGTGAAGGAATTCAACGCGCTGAGGGTGCATATATCTACATCGCACCGAACAACGGA<br>TTACTGACAAGTGTTATCGAGGAGCACGGGTACCTTGAGGCCTATGAGGTGACGAGCACAAGTGTGATCCCAGCTCGT<br>CCCGAGCCAACATTTTTCAGTCGTGAAATGGTTGCTGTGCCAGCGGCACATTTGGCAGCTGGTTTTCCTTTAAGTCAA<br>GTGGGCCGCCCTTTACAGGACTCTGAGATCGTGCGTTTTGATCGCCGCCGCCCCGCGTCGCTGGGAGACGGCGGATTC<br>GCTGGCGTGATCACGGTTGTTGACCGCCCTTACGGAAACGTATGGACGAATATTTCACGTAAAGACTTAGCGAATCAA<br>GGCATTGCATACGGAACACGTTTACGCATTACTGTGGATAATGTGCTTCCATTCGACTTGACGCTGACACCAACTTTT<br>GCTGATGCTGGTGAGATTGGAGCACCGGTGTGTTACGTAAATAGCCGTGGATACCTTTCTCTTGCACGTAACGCTAGT<br>AATTTAGCGGACACGTATAATATTCGCCGTCACATGCCAGTTAACGTCCAAGTGGTGTCCCGCACCGACGGAGAGCCG<br>CACCGTTCGGAATCATCAGAACTGGCATCGGTCAAAGGGACATAA |
| *flA* from *Peptococcaceae bacterium* (*flA*$^{Pbac}$) codon optimized for expression in *E. coli* | ATGGAAAAGGTGGACAATAAGACCGTATTTCAGAGGCCAATTATTGCGTTTATGAGTGACCTCGGCGCCTTCGATGAT<br>TCCGTAGGAATTTGCAAGGGATTAATGCTTAGCGTGTGTCCGGAAGCGCAGATCATCGACATATGTCATTCCATGACA<br>CCGTTCGACGTCGAGCAGGGGGCAAGGCTAATCGTTGATCTGCCTCGCTTTTTCCCGGAAGGAAGAACCGTATTTGCG<br>ACAACTACCTACCCTGCAACCGGTACTATGGCAAGGTCATTGGCACTACGGATTAAAAGGCCCGCGAAAGGTGGGGCA<br>CTGGGCCAATGGGCCGGAGCTGGGTTTGGTATAGAGCGCGGCGTCGGAGGTTATATATATGTAGCCCCAAACAATGGA<br>TTACTTACGGACGTCATAGAGGAACACGGATACCTAGAAGCCTATGAAATCACATCTACTAATGTGATTCCTGAAAAT<br>CCTGAGCCAACCTTTTTCAGCAGGGAGATGGTAGCCCTACCGTCTGCCTACTTAGCTGCTGGGTACCCTTTATCTGAG<br>GTTGGTCAGCCCCTAGGTGACTCAGAAATCGTAAGGTTCAAAAAGGTCCTCCCGCGAAAAATGTCGGAAAGTGAGCTC<br>GTTGGGGTCGTAGCAGCAATCGACAGGCCTTTCGGAAACGTCTGGACGAACATTTCTCGGCGAGACCTTGATAAGATT<br>GGCGTTACCTATGGGTCACAGTTAAAGGTAGTTCTCGACAATGCTTTAATGTTTGAGCTACCACTTTCGCAGACCTTT<br>GCGGATGCTAGAGAAATAGGAGCCGCTGTGGCTTATATCAACAGTCGCGGACATCTGTCCCTTGGGCGGTACGCAGCA<br>AATCTGGCCGACCGCTACAATATTAATCGTGGCATGCCAATTCGGCTAAAGGTAATTACTGGTTAA |
| *flA* from *Thermodesulforhabdus norvegica* (*flA*$^{Tnor}$) codon optimized for expression in *E. coli* | ATGGATATCGGACCCATCATAGGGTTTGCAAGTGATCTGGGCTTAAAAGACGATTCGGTTGCGCTCTGTAAAGGGTTG<br>ATGATTAGCATATGCCCGGAAGTTTATATCGTGGACATATGCCACACGATGACCCCCTTTGATATCGAGGAGGGAGCG<br>TGGTTGGCTCTTGATTTGCCCAGATTTTTTCCGGAAGGCCGAACTATATTTGCCGTTACCACATATCCTGCTACCGGC<br>ACAGAGGCTAGATCCATAGCTGTCCGAATAAAAAAAGCCGTGCCGGGTGGATCACTTGAGAAGTGGGAAGGACCGGGA<br>GGGGGTATGGAGCGTACTTTGGAGGGAGGTTACATTTATGTGGCACCGAATAATGGCTTGTTGACCTTTGTACTCGAA<br>AGATACGGGTATATTGAAGCGTATGAAATAATTTCAACTGAGTTCATTCCCGAAAATCCTGAACCTACTTTTTTCAGC<br>AGAGAAATGGTTGCAATACGGGCAGCCTGCATTGCCAAAAAAGTGGTATCACAGGGGGTGCCTCTGTCCAAAGTTGGC<br>CCTCCAATAACAGAAGATAAGCTAGCAAGAATAAAACTGTCTCTGCCGGAGAAAATAGCGCACAATGAGATAAGAGGC<br>AAGATAATTAGAATTGATTATCCTTATGGAAATGTGTGGACTAATATATCTTTCAACGATCTGAAGTCTATGGGAATT<br>AATTATGGTTCACGACTTACAGTTGTTATAGGAGATATATTGAGTTTTAATGTTTACCTGACACGGACGTTTGCCGAT<br>GCAGGCGGTATCGGAGATGTTATCTCATATATAAATAGCAGGGGTTATTTTTCTCTGGGCGGGTACGCTGCGAACCTG<br>GCCGACCTGTGTAATTTAAGAAGGGGTATGAATGTTGTTGTTATAAAGGTGTAG |

| | |
|---|---|
| *flA* from *Methanosaeta* sp. PtaU1.Bin055 (*flA*$^{\mathrm{PtaU1}}$) codon optimized for expression in *E. coli* | ATGTTGCAAAATACCAAAAATGAGTACAAACCACGCGATGGCGGAGGGCACCGCCCAATTATTGGATTCATGTCAGAT<br>TTGGGCACAGAGGACGATTCAGTGGGAATTTGTAAAGGGGTGATGTTGGGGTTATGTCCGGATGCGGTTATCGTGGAC<br>ATCAGCCATAGTATGACGCCGTGGGACATCGACCAGGGATCGCGTCTGATTGTGGACTTGCCGAAGTTCTTCCCCAAC<br>TGGACCACTTTCGCAACCACTTCCTATCGCGAAACCGGTACGTCGGCACGTTCTGTGGCTATCAAATTACCTTCTGGC<br>CACGTCTACGTTGCGCCAAATAATGGACTTTTGACACGTGTTATTGAGGACCATGGCTATGTAGAGGCGTATGAAGTG<br>ACTACAGTCGGAGCTATTCCCGCAGAGCCCGAACCGACGTGGTTTTCACGTGATATGGTCGCTTACCCTGCTGCCGCG<br>ATTGCAGCCGGCTTTCCGTTAGAGAACGTAGGTCGTCCGTTAAACGATTCCGAAATTGTACGTGCAGACTTACCCCGT<br>TATACTCAGGCTGAGGACGGCATCATTCAGGGCATCGTCACCACTATTGATCGCCCGTTCGGTAATATTTGGACTAAT<br>ATCCCACGCCGTATCATCGAAAATGAGCTTCGTATTGCCTACGGGACGAACATCCGCGTTGTACTGGATAATCTGTTA<br>GTTCTTGAAGTTCCGTTCATGCGTACCTTTGGAGACGTTGGTCTGAAGAAACCCATGTGCTATATCAACAGCCGTGGA<br>TATTTTAGCTTAGCATATTATGGTGGAAACTTAGCAGACCCCTACAATATTCGCCGTGGAATGCCGGTGAAGATCGAA<br>TCCATCACGCGTTAA |
| *flA* from *Streptomyces* sp. SAJ15 (*flA*$^{\mathrm{SAJ15}}$) codon optimized for expression in *P. putida* | ATGACGTCCAACGGGGCCCACCGCCCGATTATCGCTTTTATGAGCGATCTCGGCACGACCGACGATAGCGTGGCCCAG<br>TGTAAGGGTTTGATGCTGAGCATTTGCCCCGATGTCACGATCGTgGACGTCTGCCACAGCATGACGCCTTTTGATGTC<br>GAGGAGGGTAGCCGGTATATCGTgGACCTCCCACGTTTCTTCCCAGAAGGTACGGTGTTCGCTACGACCACGTATCCG<br>GCAACGGGGACCGAAACCCGGAGCGTCGCTGTGCGCATTCGGCAAGCAGCACAGGGTGGGGCTCGTGGTCAGTGGGCC<br>GGCTCGGGCGCAGGCTTCGAGCGGCAGGAGGGCTCCTACATTTATATCGCGCCTAACAACGGCTTGTTGACCCCGGTG<br>TTGGAAGAGCACGGCTACATCGAGGCATACGAAGTGTCCAGCACGGAGGTCATCCCAGAACGGCCAGAGCCTACCTTT<br>TACTCCCGtGAAATGGTCGCGATTCCTAGCGCACATCTCGCAGCTGGGTTTCCACTGGACCAGGTGGGTCGCCCACTG<br>AAAGACTCGGAAATTGTGCGCTTTAGCCGTCCAGCTGTCGAGACGGCTGGCACGGAGCTGACCGGGGTCGTCTCCGCA<br>ATTGATCATCCGTTCGGGAACATCTGGACGAACATTCACCGTACCGACTTGGAGAAGGCCGGGATCGGCTACGGGCGG<br>CAAATCCGGATTACGTTGGACGACGTGCTGCCTTTCGAGCTgACGCTGGTCCCTACGTTCGCGGATGCTGGCGAGGTG<br>GGTAATGTCGTCGCTTACCTGAACAGCCGCGGTTATTTGTCCCTCGCTCGCAACGCGGCGTCGTTGGCGTATCCCTAC<br>AACCTCAAGGCTGGTCTCTCGGTGCGGGTGGATGCACATTAG |
| *flA* from *Methanosaeta* sp. PtaU1.Bin055 (*flA*$^{\mathrm{PtaU1}}$) codon optimized for expression in *P. putida* | ATGTTGCAAAATACCAAAAATGAGTATAAGCCGCGCGATGGCGGGGGGCACCGCCCTATCATCGGCTTCATGTCCGAC<br>CTCGGTACGGAGGATGATAGCGTGGGCATCTGCAAGGGTGTCATGTTGGGGCTGTGTCCTGACGCCGTGATCGTGGAT<br>ATTAGCCACTCCATGACGCCCTGGGACATTGATCAAGGGTCGCGCTTGATTGTGGATCTCCCAAAGTTTTTCCCCAAC<br>TGGACGACGTTTGCGACCACGTCCTACCGCGAAACCGGCACCAGCGCACGTTCGGTCGCTATCAAGCTCCCAAGCGGG<br>CATGTCTACGTGGCGCCCAATAACGGTTTGCTGACCCGTGTGATCGAGGATCATGGGTATGTGGAGGCATACGAAGTG<br>ACCACCGTCGGCGCCATCCCAGCTGAACCGGAGCCAACGTGGTTTAGCCGTGATATGGTGGCTTACCCTGCTGCgGCT<br>ATTGCAGCGGGCTTTCCGTTGGAGAATGTGGGCCGTCCCCTGAATGATTCGGAGATTGTCCGCGCGGACCTGCCTCGG<br>TATACGCAGGCTGAAGATGGTATCATCCAGGGTATCGTGACCACCATTGATCGTCCTTTCGGTAACATTTGGACGAAT<br>ATCCCTCGGCGTATTATCGAGAACGAACTGCGGATCGCATACGGCACCAACATTCGCGTGGTCTTGGATAATTTGTTG<br>GTGCTCGAAGTGCCTTTTATGCGGACGTTTGGCGATGTCGGCCTGAAGAAACCGATGTGTTATATTAATAGCCGCGGC<br>TACTTTTCCTTGGCCTACTATGGGGGCAATTTGGCCGACCCTTACAATATTCGTCGCGGTATGCCGGTCAAAATCGAA<br>TCCATCACGCGTTAG |

**Table S5.** Nucleotide sequences of primers used for plasmid construction.

| Name | Sequence (5'→3') | Description |
|---|---|---|
| oIP001 | ATCTCTTCuGAGCACCACCACCACC | Amplification of pET-28a(+)-TEV for USER cloning, forward |
| oIP002 | ATGGCCCuGAAAATAAAGATTCTCGCCGCT | Amplification of pET-28a(+)-TEV for USER cloning with Nt His-tag, reverse |
| oIP003 | AGGGCCAuGCTACTACGAAGCGCCCAGTCC | Amplification of *flA*$^{Abar}$ for USER cloning into pET28a(+)-TEV in-phase with Nt His-tag, removes ATG, forward |
| oIP004 | AGAAGAGAuTTAAGCGACGTGCACGCGGAC | Amplification of *flA*$^{Abar}$ for USER cloning into pET28a(+)-TEV, includes TAA stop, reverse |
| oIP005 | AGGGCCAuGCATCTTCGCGTGGTAATCGC | Amplification of *flA*$_{AN11}$ for USER cloning into pET28a(+)-TEV in-phase with Nt His-tag, removes ATG, forward |
| oIP006 | AGAAGAGAuTTATCCAACCTTTACACGTACTGGAATGC | Amplification of *flA*$^{AN11}$ for USER cloning into pET28a(+)-TEV, includes TAA stop, reverse |
| oIP007 | AGGGCCAuCAAGAGGGGCAGGCCACATC | Amplification of *flA*$^{Cbac}$ for USER cloning into pET28a(+)-TEV in-phase with Nt His-tag, removes ATG, forward |
| oIP008 | AGAAGAGAuTTATGTCCCTTTGACCGATGCCAG | Amplification of *flA*$^{Cbac}$ for USER cloning into pET28a(+)-TEV, includes TAA stop, reverse |
| oIP009 | AGGGCCAuCCCGCAAACGGAAACCCTATCATC | Amplification of *flA*$^{N902}$ for USER cloning into pET28a(+)-TEV in-phase with Nt His-tag, removes ATG, forward |
| oIP010 | AGAAGAGAuTTATTTAGTCACTGTCACCGTTAAACCCG | Amplification of *flA*$^{N902}$ for USER cloning into pET28a(+)-TEV, includes TAA stop, reverse |
| oIP011 | AGGGCCAuACGACAGCTAATGGCCGTCG | Amplification of *flA*$^{Nbra1}$ for USER cloning into pET28a(+)-TEV in-phase with Nt His-tag, removes ATG, forward |
| oIP012 | AGAAGAGAuTTAACCTACCTTAACCTGCACGGAGATAC | Amplification of *flA*$^{Nbra1}$ for USER cloning into pET28a(+)-TEV, includes TAA stop, reverse |
| oIP013 | AGGGCCAuACCACGGCGAATGGACGTC | Amplification of *flA*$^{Nbra2}$ for USER cloning into pET28a(+)-TEV in-phase with Nt His-tag, removes ATG, forward |
| oIP014 | AGAAGAGAuTTAGTCAACCTTAACCTGCACTGAAATCC | Amplification of *flA*$^{Nbra2}$ for USER cloning into pET28a(+)-TEV, includes TAA stop, reverse |
| oIP015 | AGGGCCAuACTACCGCAAACGGTCGC | Amplification of *flA*$^{Nbra3}$ for USER cloning into pET28a(+)-TEV in-phase with Nt His-tag, removes ATG, forward |
| oIP016 | AGAAGAGAuTTAGCCTACTTTTACTTGAACACTAATACC | Amplification of *flA*$^{Nbra3}$ for USER cloning into pET28a(+)-TEV, includes TAA stop, reverse |
| oIP017 | AGGGCCAuTTGCAAAATACCAAAAATGAGTACAAACC | Amplification of *flA*$^{PtaU1}$ for USER cloning into pET28a(+)-TEV in-phase with Nt His-tag, removes ATG, forward |
| oIP018 | AGAAGAGAuTTAACGCGTGATGGATTCGATCTTC | Amplification of *flA*$^{PtaU1}$ for USER cloning into pET28a(+)-TEV, includes TAA stop, reverse |
| oIP019 | AGGGCCAuACGTCCAACGGTGCCCATCG | Amplification of *flA*$^{SAJ15}$ for USER cloning into pET28a(+)-TEV in-phase with Nt His-tag, removes ATG, forward |
| oIP020 | AGAAGAGAuTTAATGAGCGTCCACGCGCACC | Amplification of *flA*$^{SAJ15}$ for USER cloning into pET28a(+)-TEV, includes TAA stop, reverse |
| oIP033 | AGGGCCAUGCCGCCAACAGCACC | Amplification of *flA*$^{Scat}$ for USER cloning into pET28a(+)-TEV in-phase with Nt His-tag, removes ATG, forward |
| oIP034 | AGAAGAGAUTCAACGGGCCTCCACG | Amplification of *flA*$^{Scat}$ for USER cloning into pET28a(+)-TEV, includes TAA stop, reverse |
| oIP084 | GGAAAAA**CATATG**CATCATCACCACC | Amplification of FlA CDSs optimized for *P. putida* for cloning in pFB·1 backbone, forward, contains **NdeI** site |
| oIP085 | G**GGTACCGAGCTCGAATTC**C | Amplification of FlA CDSs optimized for *P. putida* for cloning in pFB·1 backbone, reverse, contains **EcoRI**, **SacI** and **KpnI** sites |
| oIP098 | TAGGAATTC**GAGCTC**GGTACC | Amplification of pFB·1 for production of His-tag fusions, forward, contains **SacI** site |
| oIP099 | TGATGATG**CATATG**TTTTTCCTCC | Amplification of pFB·1 for production of His-tag fusions, forward, contains **NdeI** site |
| pET_flA_rv | ATGGCCCUGAAAATAAAGATTCTCGCCGC | Amplification of pET-28a(+)-TEV for USER cloning, forward |
| pET_flA_fw | AGGCCCGTUGATCGAGCACCACCACCACC | Amplification of pET-28a(+)-TEV for USER cloning with Nt His-tag, reverse |
| flA_fw_USER | AGGGCCAUGCCGCCAACGGCAGCCAG | Amplification of *flA*$^{MA37}$ for USER cloning into pET28a(+)-TEV in-phase with Nt His-tag, removes ATG, forward |
| flA_rev_USER | AACGGGCCUCCACGCGCACCTTCAGGCC | Amplification of *flA*$^{MA37}$ for USER cloning into pET28a(+)-TEV, includes TGA stop, reverse |
| Fw2 FlA1 A0A1I4QPY6 | AGGGCCAUATGGATATCGGACCCATC | Amplification of *flA*$^{Tnor}$ for USER cloning into pET28a(+)-TEV in-phase with Nt His-tag, removes ATG, forward |
| Rv FlA1 A0A1I4QPY6 | AGAAGAGAUCTACACCTTTATAACAACAAC | Amplification of *flA*$^{Tnor}$ for USER cloning into pET28a(+)-TEV, includes TGA stop, reverse |
| Fw FlA1 A0A1G9FQX8 | AGGGCCAUTCGGACAGTTATTCGCG | Amplification of *flA*$^{Amza}$ for USER cloning into pET28a(+)-TEV in-phase with Nt His-tag, removes ATG, forward |

| | | |
|---|---|---|
| Rv FlA1 A0A1G9FQX8 | AGAAGAGAUCTACGACCGGGTCACTCTGAC | Amplification of *flA*$^{Amza}$ for USER cloning into pET28a(+)-TEV, includes TGA stop, reverse |
| Fw FlA1 A0A0J1FI89 | AGGGCCAUGAAAAGGTGGACAATAAG | Amplification of *flA*$^{Pbac}$ for USER cloning into pET28a(+)-TEV in-phase with Nt His-tag, removes ATG, forward |
| Rv FlA1 A0A0J1FI89 | AGAAGAGAUTTAACCAGTAATTACCTTTAG CCG | Amplification of *flA*$^{Pbac}$ for USER cloning into pET28a(+)-TEV, includes TGA stop, reverse |
| Fw FlA1 UPI000CD15D3E | AGGGCCAUGCGGTCACCGTCG | Amplification of *flA*$^{CA12}$ for USER cloning into pET28a(+)-TEV in-phase with Nt His-tag, removes ATG, forward |
| Rv FlA1 UPI000CD15D3E | AGAAGAGAUCGCCGAGGAAGGACAGGG | Amplification of *flA*$^{CA12}$ for USER cloning into pET28a(+)-TEV, includes TGA stop, reverse |

**Figure S1.** Phylogenetic tree of 16S rRNA sequences from organisms encoding known and putative fluorinases.

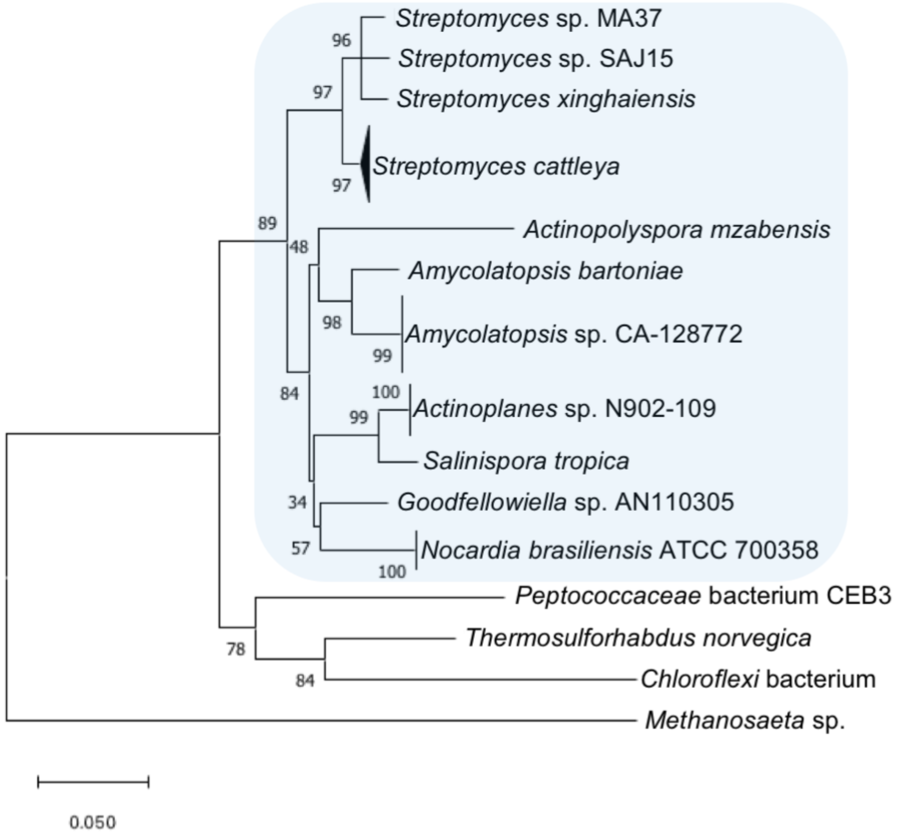


Searches were performed using the NCBI BLASTn tool against the selected organisms, using the 16S rRNA sequence from *Streptomyces* sp. MA37 as query. Sequence accession numbers are presented in **Table S2**. In the case of *Nocardia brasiliensis*, sequences from strain ATCC 700358 were retrieved. Sequences specific to *Methanosaeta* sp. PtaU1.Bin055 could not be retrieved as the whole genome sequence is not available, and the 16S rRNA sequence from a closely related *Methanosaeta* sp. was used instead. When more than one sequence were obtained from the whole-genome sequence of the same organism, these were all included for multiple sequence alignment and phylogenetic tree construction using the MEGA software (sub-trees collapsed for clarity). The phylogenetic tree was inferred using the Neighbor-Joining method with a bootstrap of 10,000 iterations. The percentage of replicate trees in which the associated taxa clustered together in the boot-strap test are shown next to the branches. The tree is drawn to scale, with branch lengths in the same units as those of the evolutionary distances used to infer the phylogenetic tree. Sequences sourced from Actinomycetes are highlighted with a light blue square.

**Figure S2.** SDS-PAGE evaluation of FIAs produced in recombinant *E. coli* BL21(DE3).

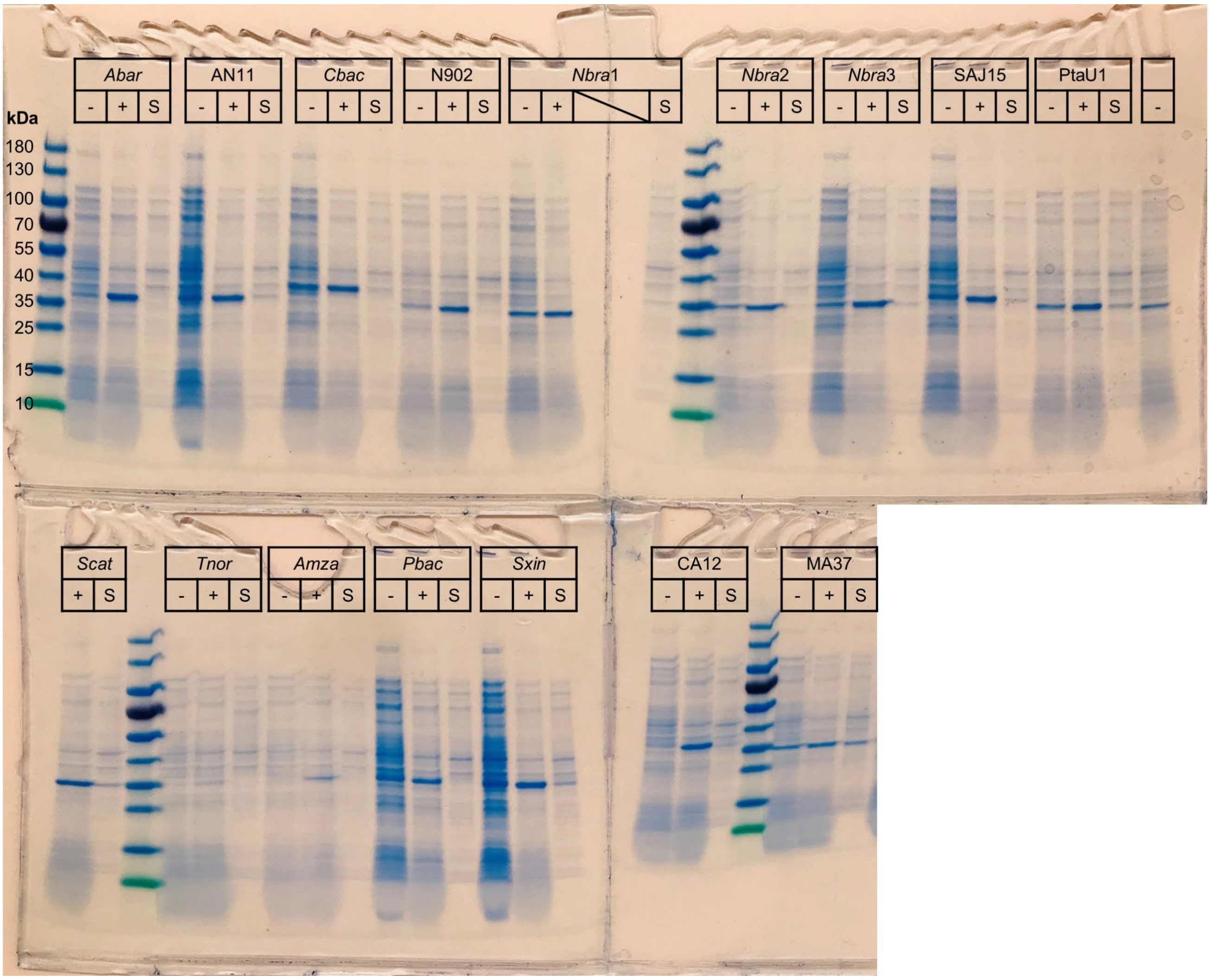


Cultures were carried out in 96-well plates. Lane labels indicate whole-cell extracts from non-induced samples (–), whole-cell extracts from induced samples (+) and soluble fraction from induced samples (S).

**Figure S3.** Preliminary activity assays of putative fluorinases at increasing SAM concentrations.

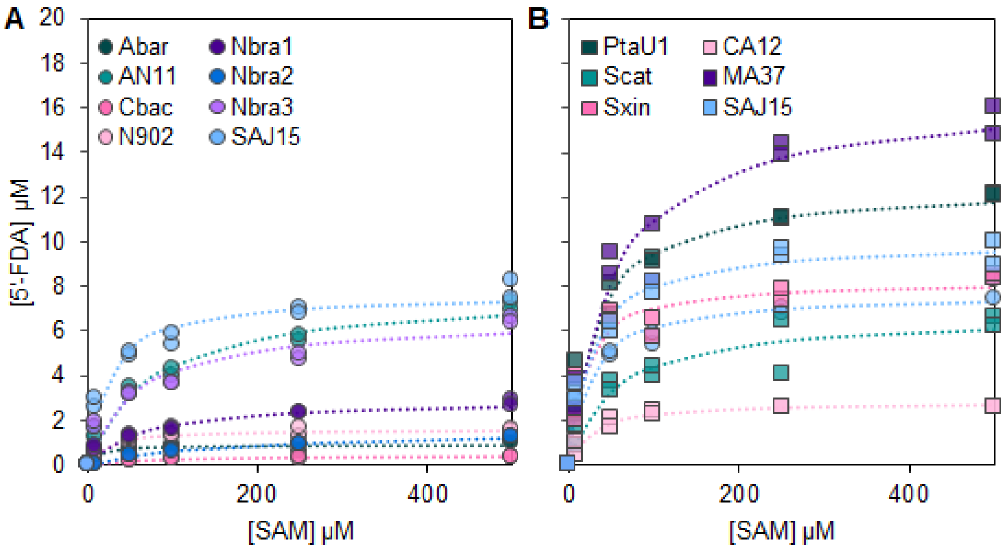


Symbols indicate the 5'-fluoro-5'-deoxyadenosine (5'-FDA) concentration for individual replicates. Dotted lines represent fit to Michaelis-Menten equation. Panels **A** and **B** show average data from individual experiments carried out in different days in triplicates. Data for the fluorinase of strain SAJ15 from panel **A** (light blue circles) is also shown in panel **B** to facilitate comparison. The reactions were carried out at 37°C in 50 mM HEPES buffer, pH = 7.8, if the presence of 75 mM KF.

**Figure S4.** Residue conservation for putative fluorinases/chlorinases retrieved from the EnzymeMiner search.

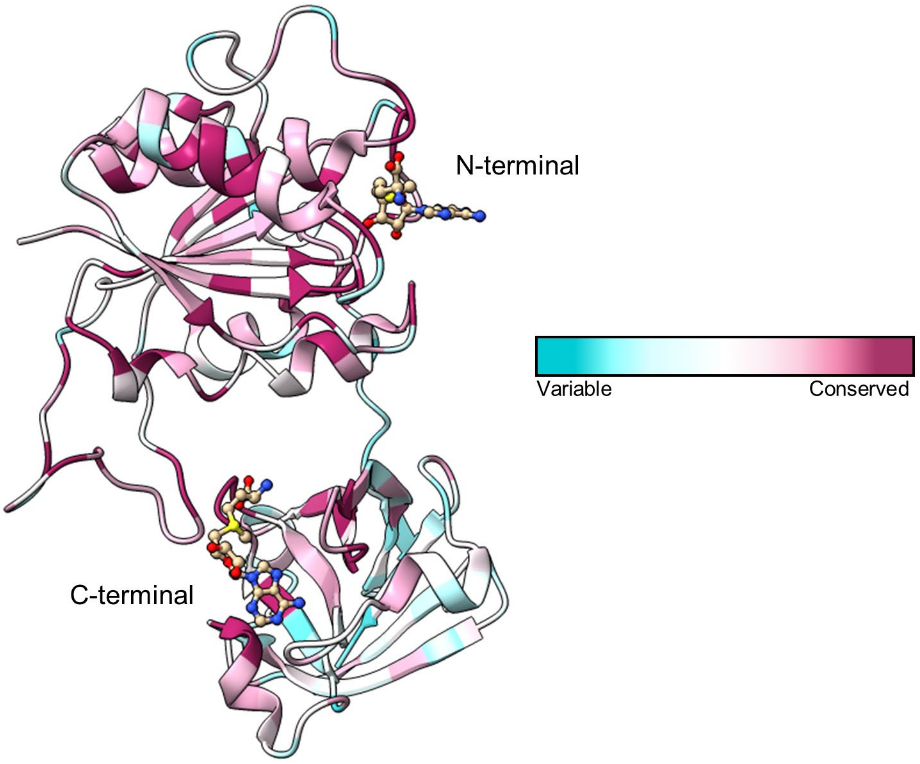


The degree of conservation for each amino acid position in fluorinase/chlorinase enzymes was estimated using the ConSurf server and the results were visualized with ChimeraX. A single fluorinase monomer, based on the crystal structure of FlA$^{Scat}$ (PDB 1RQP), is shown as a cartoon representation. Two SAM substrate molecules, bound to the *N*-terminal and *C*-terminal domains, respectively, are shown in ball-and-stick representation.
